# Dietary L-3,4-dihydroxyphenylalanine (L-DOPA) augments cuticular melanization in *Anopheles* mosquitos while reducing their lifespan and malaria parasite burden

**DOI:** 10.1101/2024.09.30.615839

**Authors:** Emma Camacho, Yuemei Dong, Christine Chrissian, Radames J.B. Cordero, Raúl G. Saraiva, Yessenia Anglero-Rodriguez, Daniel F.Q. Smith, Ella Jacobs, Isabelle Hartshorn, Jose Alberto Patiño-Medina, Michael DePasquale, Amanda Dziedzic, Anne Jedlicka, Barbara Smith, Godfree Mlambo, Abhai Tripathi, Nichole A. Broderick, Ruth E. Stark, George Dimopoulos, Arturo Casadevall

**Affiliations:** Johns Hopkins University, Baltimore, USA; City College of New York, New York, USA

## Abstract

L-3,4-dihydroxyphenylalanine (L-DOPA), a naturally occurring tyrosine derivative, is prevalent in environments that include mosquito habitats, potentially serving as part of their diet. Given its role as a precursor for melanin synthesis we investigate the effect of dietary L-DOPA on mosquito physiology and immunity to *Plasmodium falciparum* and *Cryptococcus neoformans* infection. Dietary L-DOPA is incorporated into mosquito melanin via a non-canonical pathway and has a profound transcriptional effect associated with enhanced immunity, increased pigmentation, and reduced lifespan. Increased melanization results in an enhanced capacity to absorb electromagnetic radiation that affects mosquito temperatures. Bacteria in the mosquito microbiome act as sources of dopamine, a substrate for melanization. Our results illustrate how an environmentally abundant amino acid analogue can affect mosquito physiology and suggest its potential usefulness as an environmentally friendly vector control agent to reduce malaria transmission, warranting further research and field studies.

## INTRODUCTION

Malaria is a major public health threat exacerbated by the impact of climate change ^1^. Global warming has pushed mosquitoes to new latitudes and altitudes, prolonging the malaria transmission season and increasing the risk to previously unaffected communities ^2,3^. In addition, human activities such as land development, intensive agricultural practices, and the widespread use of chemical pesticides significantly influence the proliferation and spread of mosquito populations ^2^. As a result, the dynamics of disease transmission by these vectors are continually evolving. A stark illustration of these public health consequences occurred last year when the United States of America (USA) experienced nine cases of locally acquired malaria –marking the first time in two decades that locally acquired transmission had occurred. Notably, eight of these cases were reported in Florida and Texas, with an additional case identified in Maryland. This alarming trend contrasts with the situation in 2003 when eight cases of locally acquired malaria were identified and all were restricted to Florida’s Palm Beach County. The *Anopheles* mosquito vectors, responsible for transmitting malaria, are found throughout many regions of the US, posing a substantial risk if they encounter a malaria-infected person ^4^. In Africa, climate change is fueling a perfect storm for malaria ^3,5^. The combination of increased frequency of extreme weather events, deforestation, warmer temperatures at higher altitudes, rapid spread of insecticide-resistant *Anopheles* mosquitoes ^6^, and emergence of urban mosquito vectors like *Anopheles stephensi* ^7^ threaten progress in malaria control and pose greater risks to human health.

Most vector borne diseases have been mitigated through the implementation of efficacious vector control programs ^8^. However, maintaining the long-term sustainability of such programs is difficult and often impractical. In the case of malaria, prior to the use of insecticides, vector control strategies relied on understanding the ecology of local vectors such as changing the aquatic habitats where the vector developed (e.g., draining swamps, applying oil to open water bodies, cutting vegetation at edges of ponds/streams). These interventions were highly successful but labor intensive and they had huge negative environmental impact ^9^. Consequently, environmental management practices were replaced by insecticide-based vector control. Despite initial success – between 2000 to 2015, the employment of nets impregnated with residual insecticides led to a reduction in over 50% of malaria cases in Africa – these efforts were compromised by the emergence of insecticide-resistance in the mosquito vectors and drug-resistance in the malaria- carrying parasites which they harbor ^10,11^.

Considering the challenges outlined above, the World Health Organization (WHO) has urged the development of cost-effective, sustainable, and environmentally safe vector-control interventions. As adult mosquitoes of both sexes, including hematophagous species, feed from floral nectar as a common nutritional source ^12^, attractive toxic sugar baits (ATSBs) are alternative and potential new tools ^13^. These mosquito control methods rely on appealing cues and sugar loaded with distinct agents such as transgenic entomopathogenic fungi ^14^, genetically modified bacteria (para- transgenesis) that confer mosquito refractoriness ^15^, or natural microbiota that result in increased resistance and/or decreased fitness ^16^.

In insects, melanin plays an essential role in three physiological processes: cuticular melanization, wound healing, and immunity ^17^. Research on the complex nature of plant-insect interactions has shown that insects adapt to L-DOPA-rich diets ^18–20^. For instance, the pea aphid *Acyrthosiphon pisum* uses plant-derived L-DOPA to promote melanin synthesis for wound healing and UV protection while sequestering high toxic compound levels ^18^. Consequently, we aim to investigate the potential of 3,4-dihydroxyphenylalanine (L-DOPA), a melanin precursor found naturally in flowers of the faba bean, *Vicia faba* L. (Fabaceae) ^21,22^. Extrafloral nectaries of these legumes also emit headspace volatiles like linalool, known to attract *Culex*, *Aedes* and *Anopheles* mosquitoes ^23,24^. We hypothesize that adult *A. gambiae* mosquitoes modulate melanin-related physiological processes in response to a sugar diet supplemented with L-DOPA, thereby reducing their olfactory susceptibility to foreign organisms that could pose a threat to humans. Consequently, we assess the oral toxicity of L-DOPA concentrations in a sugar solution, define the transcriptional profile of *A. gambiae* female mosquitoes in response to dietary L-DOPA and evaluate its functional impact on key vector life traits (Fig. 1). L-DOPA-fed *A. gambiae* female mosquitoes exhibit downregulation of essential immune-related genes, have reduced malaria parasite and fungal burden, enhanced cuticular melanization, shortened life span, and suppress expression of critical genes involved in the synthesis of cuticular hydrocarbons. Our findings suggest a potential role for L-DOPA in malaria control strategies that target mosquito reproduction.

**Fig 1.**
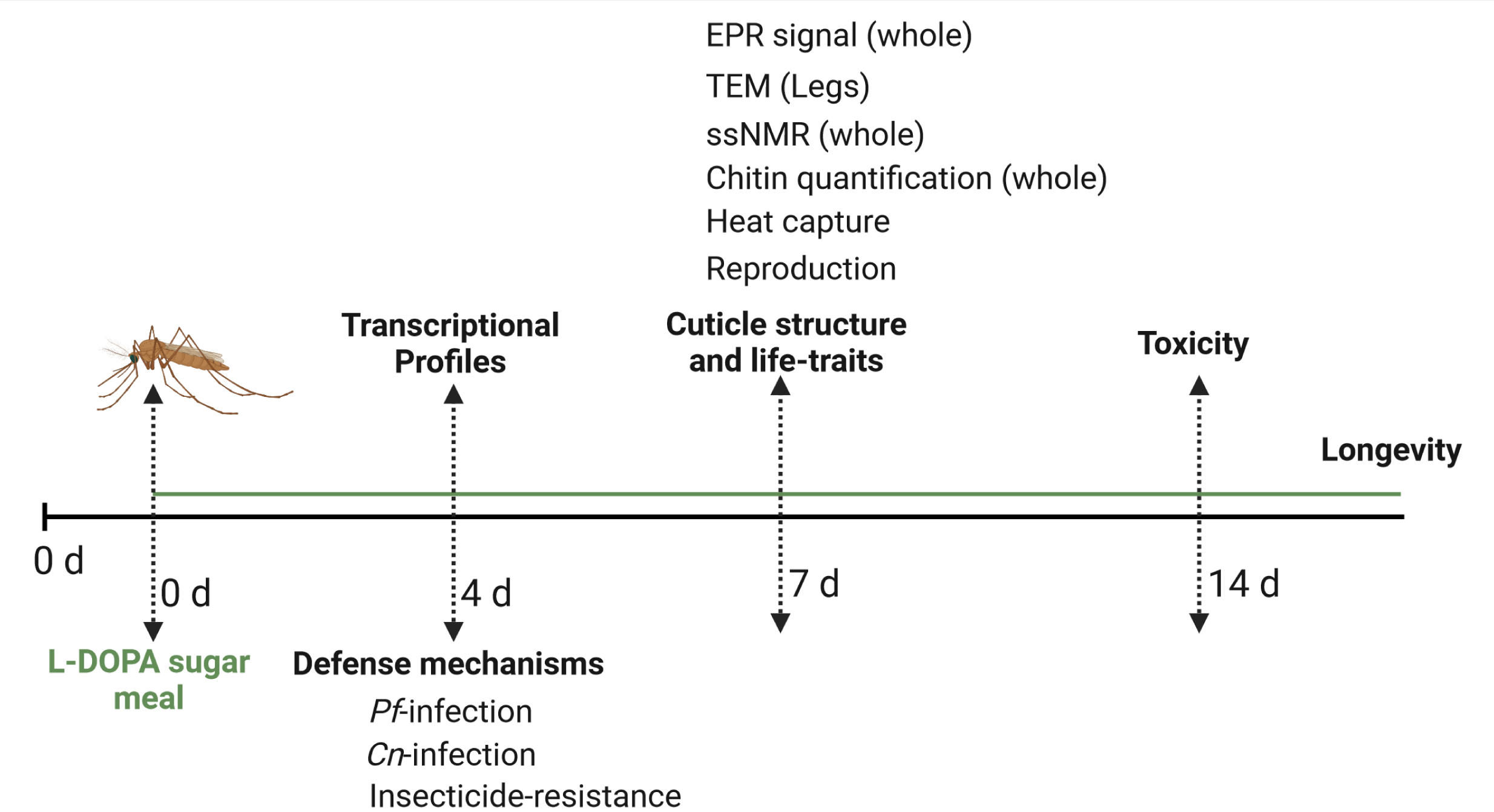
Experimental overview for investigating the impact of L-DOPA-supplemented sugar meal on *A. gambiae* mosquitoes. Created in BioRender. Camacho, E. (2025) https://BioRender.com/a17m623

## RESULTS

### Effects of dietary L-DOPA on survival of *A. gambiae* female mosquitoes

*A. gambiae* females rely mainly on a blood meal to obtain phenylalanine as the initial substrate for the melanization process. In contrast, male mosquitoes depend exclusively on plant sources (Fig. 2). To investigate whether a sugar solution supplemented with L-DOPA had a beneficial or detrimental impact on melanin-related physiological processes of adult mosquitoes, we first assessed its effect as a supplement to sugar meals on longevity, as a measure of mosquito fitness. We provided 3–4-day-old female mosquitoes with a low sugar solution supplemented with L-DOPA at concentrations ranging from 0.2 to 10 mM (0.4 to 20% w/w) and then monitored their survival over a 14-d period. These L-DOPA concentrations were comparable to those reported in natural sources, which range from 0.2 to 7.7% (w/w) ^21,25–30^. We observed that after 8 d of L- DOPA exposure, mosquitoes fed on L-DOPA solutions at most concentrations started to experience a drastic increase in mortality, except for those fed on 1 mM (2%) L-DOPA, whose survival exceeded that of the control (Fig. 3a). By 14 d, mosquitoes fed with L-DOPA solutions at concentrations lower than 1 mM exhibited a mortality of ∼50%, while at higher doses mortality increased in a dose-response relationship. Notably, analysis of the accumulative survival at different time points revealed that dietary L-DOPA exhibited a clear biphasic dose-response (Fig. 3b). Conversely, when providing dopamine in the sugar meal the hormesis effect disappeared, with control mosquitoes and those fed with up to 1 mM dopamine-sugar dying at an equal pace (Figs. 3c, 3d). Consequently, we used 1 mM L-DOPA in subsequent experiments.

**Fig 2.**
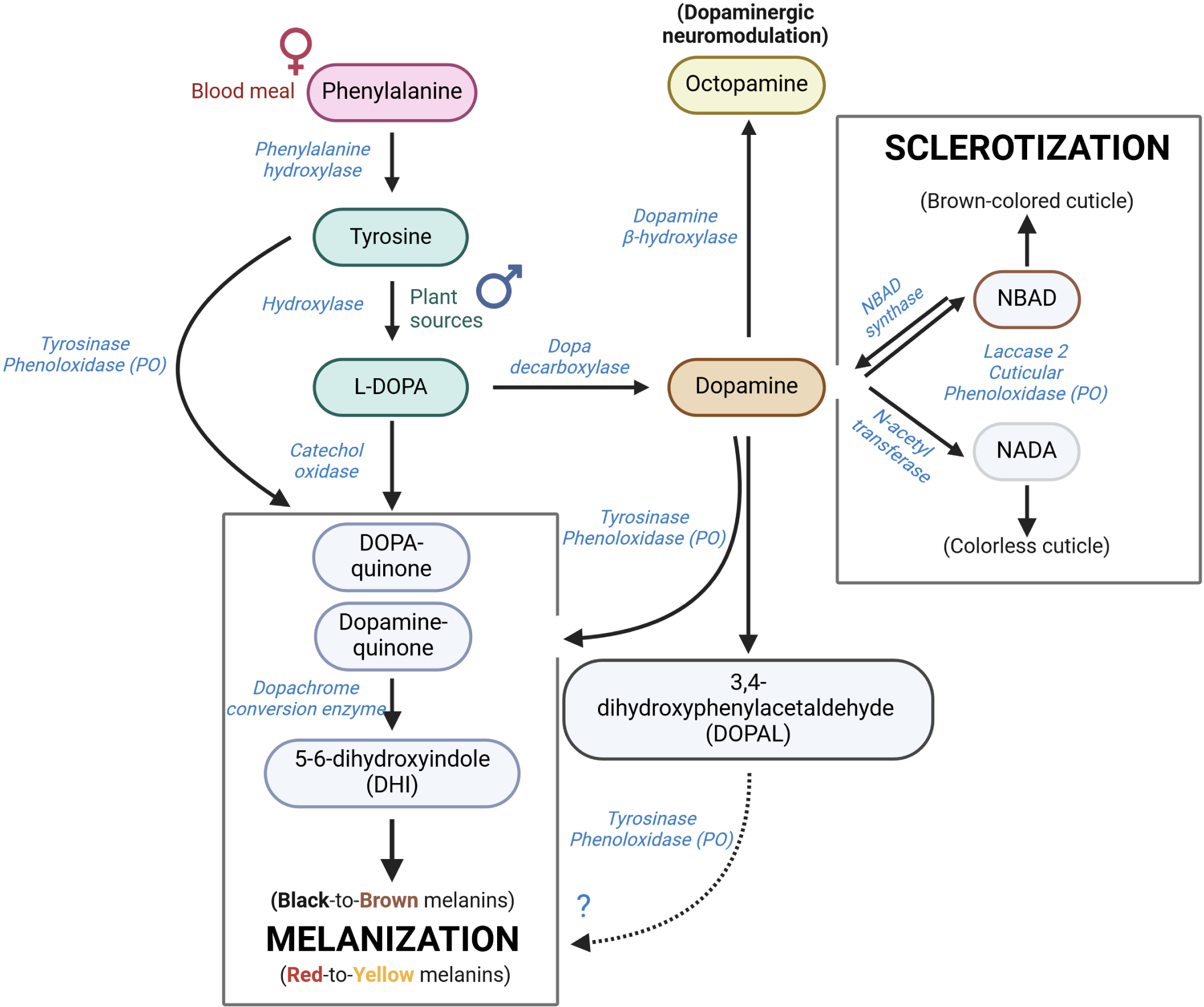
Melanization pathways in insects. In *Anopheles* female mosquitoes, melanin synthesis relies mostly on dopamine produced from the blood meal-derived phenylalanine; however, other precursors from plant sources such as tyrosine or L-DOPA are used by both female and male mosquitoes for the generation of this biopolymer. Most melanins synthesized in insects derive from L-DOPA or dopamine in a reaction initiated by phenoloxidases present in the hemolymph; however, there is one possible minor pathway from a degradation product of dopamine: 3,4- dihydroxyphenylaldehide (DOPAL). DOPAL is a highly unstable compound that reacts with primary amines, leading to protein crosslinking and inactivation. Current literature suggests that its toxicity outweighs its benefits; evidence for DOPAL melanin formation in any biological system remains to be established. Simultaneously but via an independent pathway, dopamine serves as substrate for a process known as sclerotization which promotes hardening of insect cuticles (exoskeleton) and prevents desiccation, dehydration, and invasion by foreign organisms. This process is catalyzed by cuticle-bound phenoloxidases ^12,17^. Created in BioRender. Camacho, E. (2025) https://BioRender.com/d90z642

**Fig 3.**
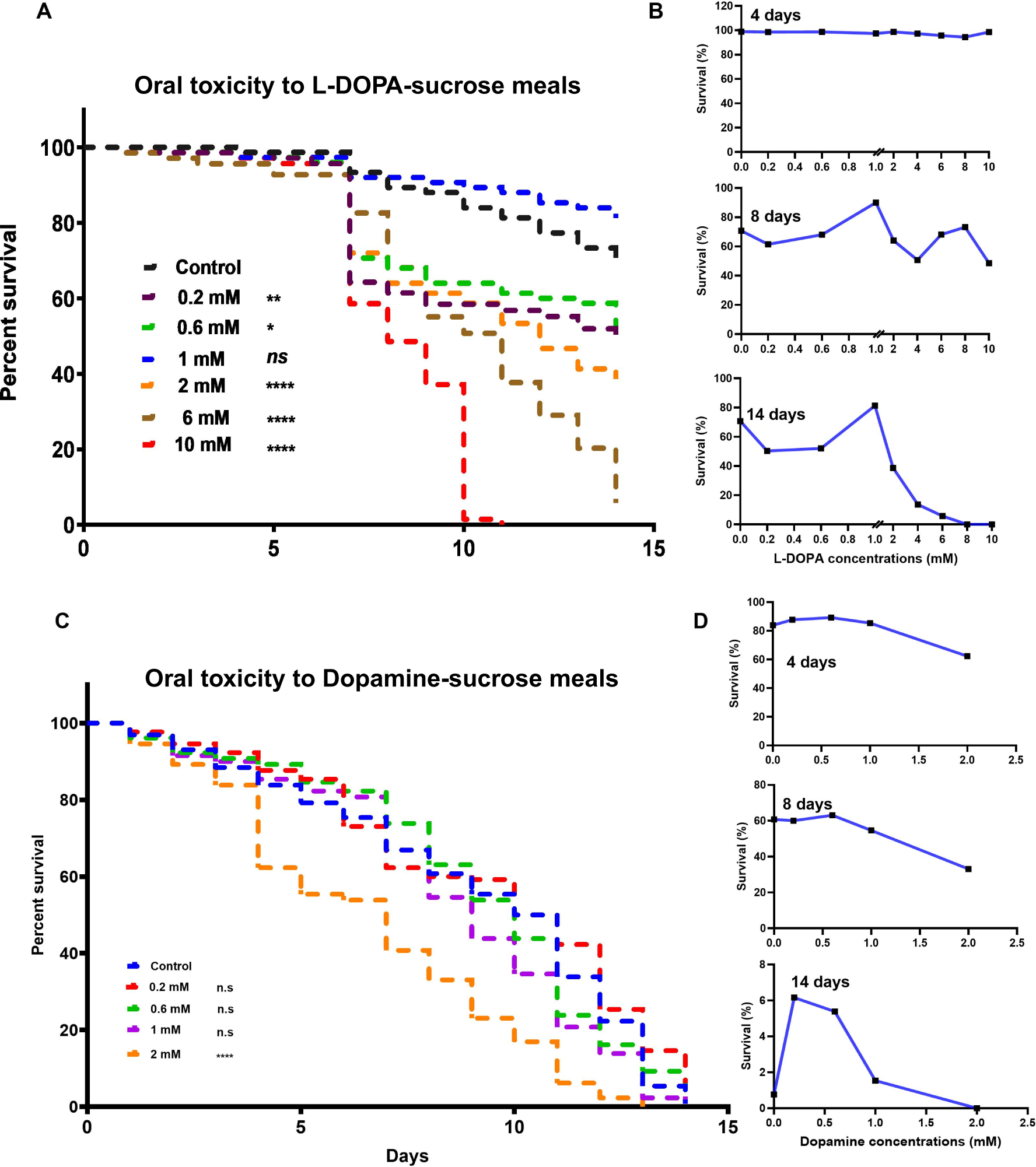
L-DOPA supplementation exerts a hormesis effect. **a** Daily proportion of surviving mosquitoes after L-DOPA feeding in 1.5% sugar meal at various supplementation concentrations. Each survival curve represents pooled data from three independent experiments (n= 75). Survival curves for L-DOPA-fed mosquitoes were compared statistically to the survival curves of control mosquitoes using Log-rank Mantel-Cox tests; ns not significant, \**p =* 0.0113 0.6 mM, \*\**p =* 0.0007 0.2 mM, \*\*\*\**p <* 0.0001 2 mM - 10 mM. **b** Cumulative survival percentage at days 4, 8 and 14 shows how mosquitoes adapt to the toxic effects of L-DOPA at concentrations below and above 1 mM over time. **c** Daily proportion of surviving mosquitoes after dopamine feeding in 1.5% sugar meal at various supplementation concentrations. Each survival curve represents pooled data from three independent experiments (n= 75). Survival curves for dopamine-fed mosquitoes were compared statistically to the survival curves of control mosquitoes using Log-rank Mantel-Cox tests; ns not significant, \*\*\*\**p <* 0.0001 2 mM. **d** Cumulative survival percentage at days 4, 8 and 14 shows highly toxic effects of dopamine over time and striking death observations for sugar-fed control mosquitoes.

### Dietary L-DOPA affects expression of mosquito immune- and cuticle-related genes

To identify genes whose expression was affected by a non-toxic concentration of L-DOPA, we collected female *A. gambiae* mosquitoes post 1 mM L-DOPA feeding of the sugar meal during 4 d and performed RNA sequencing (mRNA-seq) on whole mosquitoes (headless) and midguts. We observed 902 and 165 differentially expressed genes between L-DOPA-fed versus sucrose- fed controls (≥ 2-fold change, *P* < 0.05) in the whole mosquitoes and midgut samples, respectively (Fig. 4). In whole mosquitoes, these included 808 downregulated and 94 upregulated genes, both overrepresented in the functional category of stress-related (R/S/M; oxidoreductive, stress- related, mitochondrial) (Figs. 4a, 4b). In midguts, we observed 50 downregulated and 115 upregulated genes. Downregulated genes were overrepresented in the functional categories of transport, DNA-related (R/T/T; replication, transcription, translation), and stress-related (Figs. 4c, 4d, Supplementary Data 1, Supplementary Fig. 1). Unexpectedly, key genes involved in the melanin biosynthesis pathway such as prophenoloxidase 5 (PPO5) (AGAP012616) and dopachrome-conversion enzyme (DCE) (AGAP005959) were significantly downregulated. Diverse immune-related gene families involved in pattern recognition receptors (PRRs) (e.g., *peptidoglycan recognition proteins* (PGRP), *scavenger receptor* (SCR), and *galectins* (GALE)), signal modulation and amplification (e.g., *clip domain serine proteases* (CLIP) and *serpins* (SRPN)), signal transduction pathways (e.g., TOLL), and other effector systems (e.g., *caspases* (CASP) also exhibited significant downregulation. Furthermore, two critical genes for dopamine metabolism in insects, dopamine beta-monooxygenase (DBH) (AGAP010485) and dopa decarboxylase (DDC) (AGAP009091), which participate in the synthesis of octopamine (main neurotransmitter) and cuticle, respectively, were also downregulated. By contrast, the immune- related dally protein (AGAP005799) involved in signal transduction was significantly upregulated (Fig. 5a, upper panel, Supplementary Data 1). In the midguts, the most significantly upregulated gene corresponded to phosphoinositide-3-kinase, regulatory subunit (PI3K) (AGAP005583) that participates in bacterial phagocytosis and apoptosis, and autophagy in *Drosophila*^31^ (Fig. 5a, lower panel; Supplementary Data 1).

**Fig 4.**
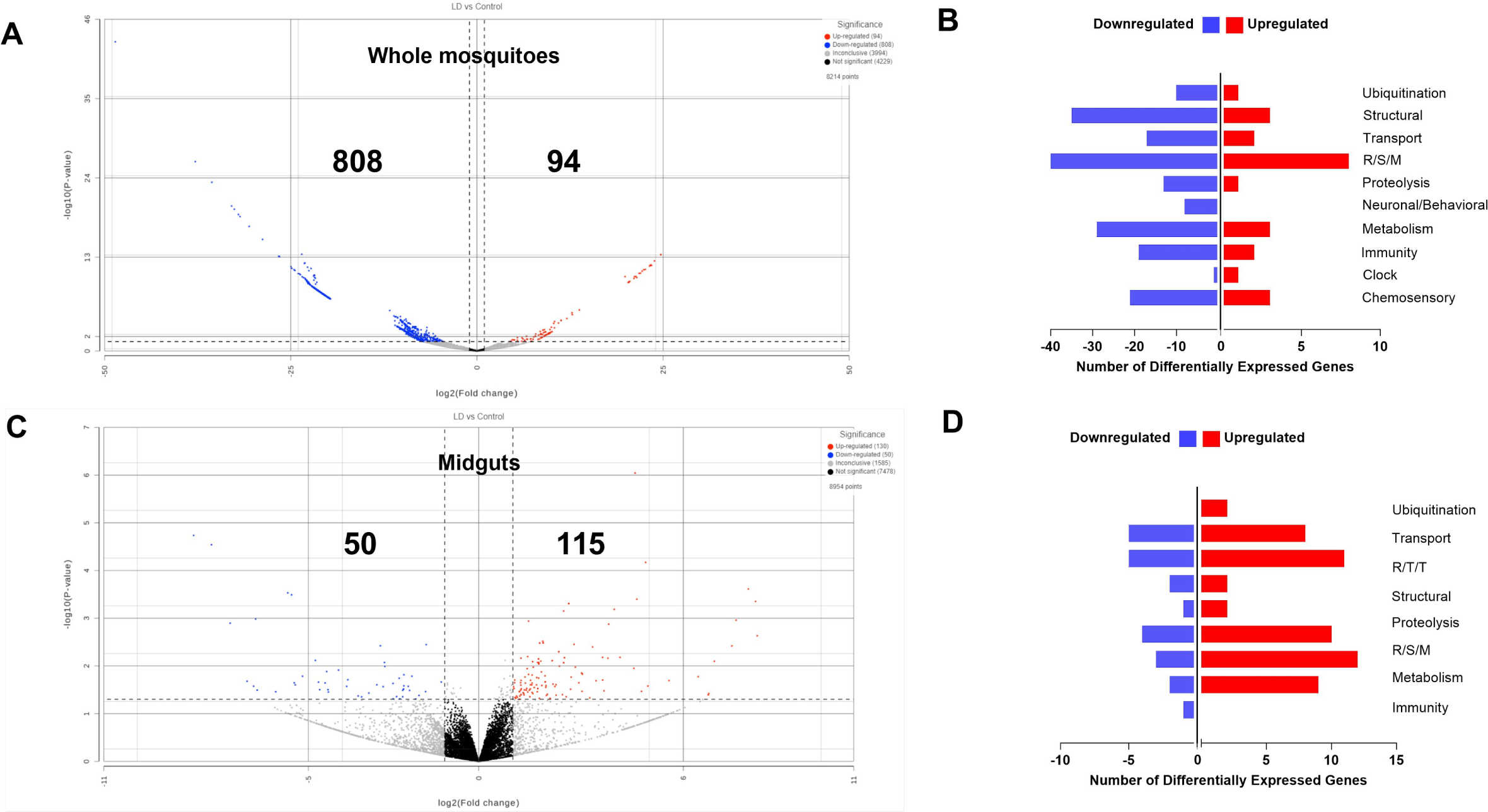
Comparative transcriptome analysis of *A. gambiae* gene expression for L-DOPA-fed and sugar-fed female mosquitoes. **a** Volcano plot of differentially expressed genes (DEGs) in whole (headless) mosquitoes. Significantly upregulated and downregulated genes are highlighted in red and blue spots, respectively. **b** DEGs by gene ontology category in whole L-DOPA-fed female mosquitoes compared with sugar-fed (control) females. Differential gene expression was considered significant when the expression fold change (L-DOPA-fed versus sugar-fed) was ≥1.3 on a -log10 scale (P < 0.05). Categories are R/S/M, oxidoreductive, stress-related, and mitochondrial. **c** Volcano plot of differentially expressed genes (DEGs) in mosquito midguts. Significantly upregulated and downregulated genes are highlighted in red and blue spots, respectively. **d** DEGs by gene ontology category in midguts from L-DOPA-fed female mosquitoes compared with sugar-fed females. Differential gene expression was considered significant when the expression fold change (L-DOPA-fed versus sugar-fed) was ≥1.3 on a -log10 scale (P < 0.05). Categories are R/T/T, replication, transcription, and translation; R/S/M, oxidoreductive, stress- related, and mitochondrial.

**Fig 5.**
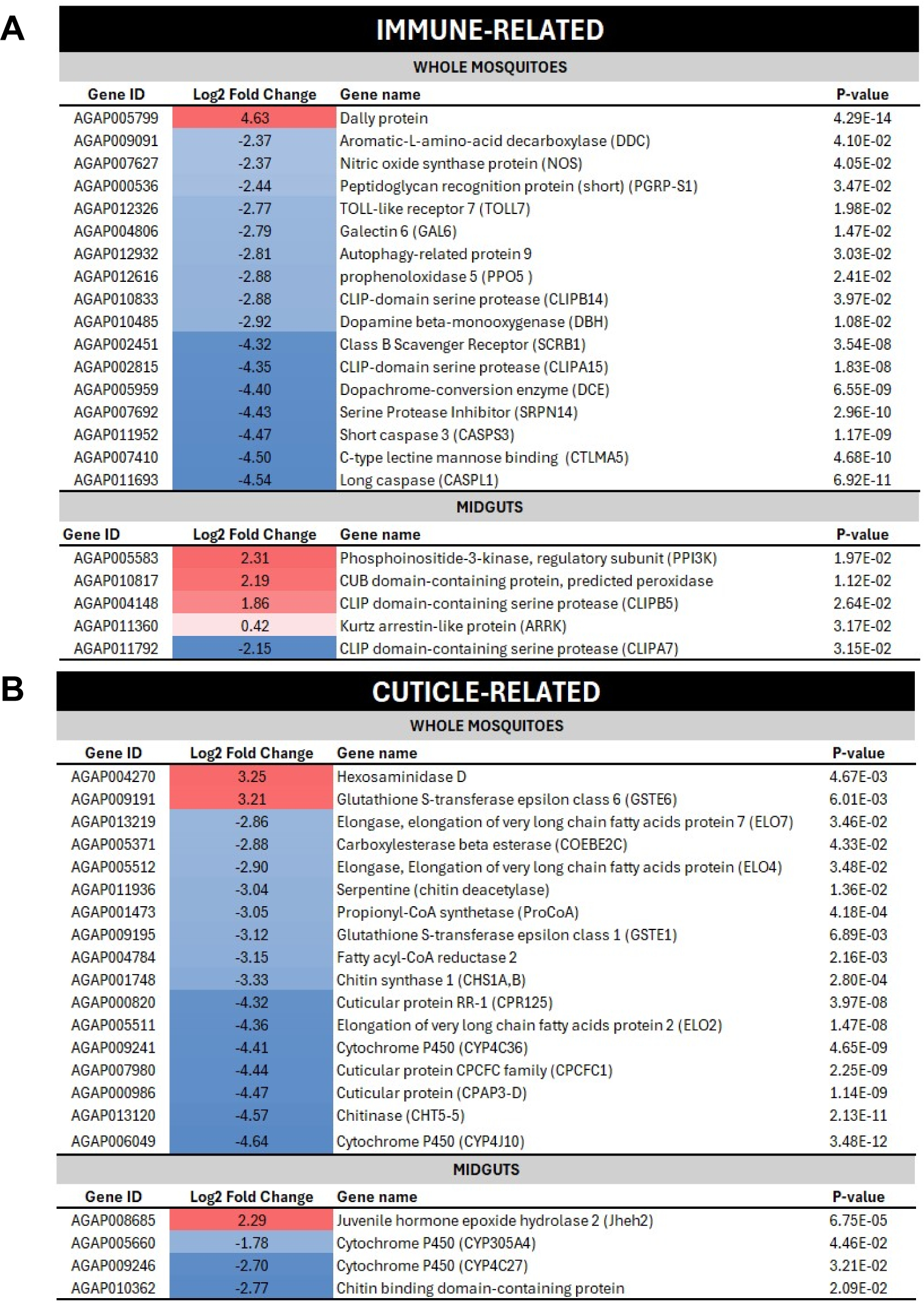
Differential transcript abundances of melanin-related physiological processes in *A. gambiae* female mosquitoes following feeding of a sugar-meal supplemented with 1 mM L- DOPA for 4 d. Log2 scale indicates transcripts that were found at significantly higher (red) or lower (blue) levels in whole (headless) mosquitoes (upper panel) and midguts (lower panel) from*A. gambiae* post L-DOPA-feeding versus sugar-fed (controls). **a** Set of immune-related transcripts. **b** Set of cuticle-related transcripts.

Regarding cuticle-related genes, transcriptional profiles of whole mosquitoes showed that L- DOPA-supplementation was associated with an extensive downregulation of gene families involved in cuticle regulation, metabolic detoxification, and synthesis of cuticular hydrocarbons. These included: cuticular proteins (e.g., AGAP000820 and AGAP000986); chitinase (CHT5-5, AGAP013120); chitin synthase 1 (CHS1A, AGAP001748); cytochrome P450s; propionyl-CoA- synthase (AGAP001473); and elongases (AGAP005511, AGAP005512, AGAP013219). Notably, upregulated genes in L-DOPA-fed mosquitoes corresponded to hexosaminidase D (AGAP004270) and glutathione S-transferase epsilon class 6 (GSTE6) (AGAP009191) (Fig. 5b, upper panel, Supplementary Data 1). The latter gene is associated with the development of insecticide resistance in the malaria vector *Anopheles funestus* ^32^. Similarly, in the midgut, cytochrome P450s and chitin-related genes were downregulated while *Jheh2*, a gene coding for juvenile hormone epoxide hydrolase, was upregulated (Fig. 5b, upper panel, Supplementary Data 1).

Notably, almost a third (255 out of 902) of the genes differentially regulated in response to L- DOPA corresponded to hemocyte-associated genes. Most of these were oenocytoid-related (76.9%) versus granulocyte-related immune cell subtypes (19.2%), as defined by scRNA-seq hemocytes clustering performed by Kwon *et al* ^33^. These genes are involved in multiple functional categories including immunity, lipid metabolism, proteolysis, transport, and chemosensory systems such as odorant binding proteins (OBPs) and G-coupled protein receptors (GPCRs) (Supplementary Data 1). Collectively, our transcriptional analysis suggests that dietary L-DOPA was associated with a global modulation of genes beyond those related to melanin-associated processes.

### Dietary L-DOPA promotes mosquito resistance to malaria parasite and fungal infection

To investigate whether a L-DOPA-supplemented sugar meal could modulate *P. falciparum* infection in *A. gambiae* mosquitoes, we fed *Anopheles* female mosquitoes on L-DOPA containing solutions (0.2 mM, 0.6 mM, and 1 mM) for 4 d prior to *P. falciparum* infection. The intensity of the parasite infection was assessed by enumeration of *P. falciparum* oocysts at 8 d post-infection (dpi). Mosquitoes fed with a sugar meal supplemented with L-DOPA demonstrated a dose- dependent reduction of the *P. falciparum* parasite burden with a significant reduction for the 1 mM L-DOPA in comparison to the sugar-fed control (median number of oocysts 2 vs 5, *p*= 0.0081). L- DOPA supplementation also reduced the infection prevalence (77.5% vs 87.3% at 1 mM in comparison to the control) (Fig. 6a, left panel). Using dopamine instead of L-DOPA in the sugar meal was also associated with a significant reduction of *P. falciparum* infection burden in a dose- dependent manner relative to the sugar-fed controls. At the highest dose, 1 mM dopamine, the median number of oocysts was 2 vs 7, *p*=0.0019 in comparison to the control, and the prevalence of infection was also reduced (81.1% vs 87.5%) but to a lesser degree than in the presence of an L-DOPA-supplemented sugar meal (Fig. 6a, right panel). No melanized parasites were observed in these experiments at either 2 or 8 dpi, regardless of whether L-DOPA- or dopamine- supplemented sugar meals were provided following parasite infection (data not shown), indicating that the reduction in parasite infection was due to a non-melanin-based immune response. Given that nectars provide mosquitoes with L-DOPA, we investigated whether L-DOPA directly exhibits *in vitro* antiplasmodial activity to both sexual and asexual stages of the human malaria parasite. At 1 mM L-DOPA, sexual stages of *P. falciparum* exhibited significant toxicity (32.4% survival vs sugar-fed control, *p* < 0.0001), while the viability of asexual stages of the parasite was nearly abolished at 1 mM L-DOPA and drastically reduced at 0.5 mM L-DOPA (26.3% survival vs control) (Fig. 6b).

**Fig 6.**
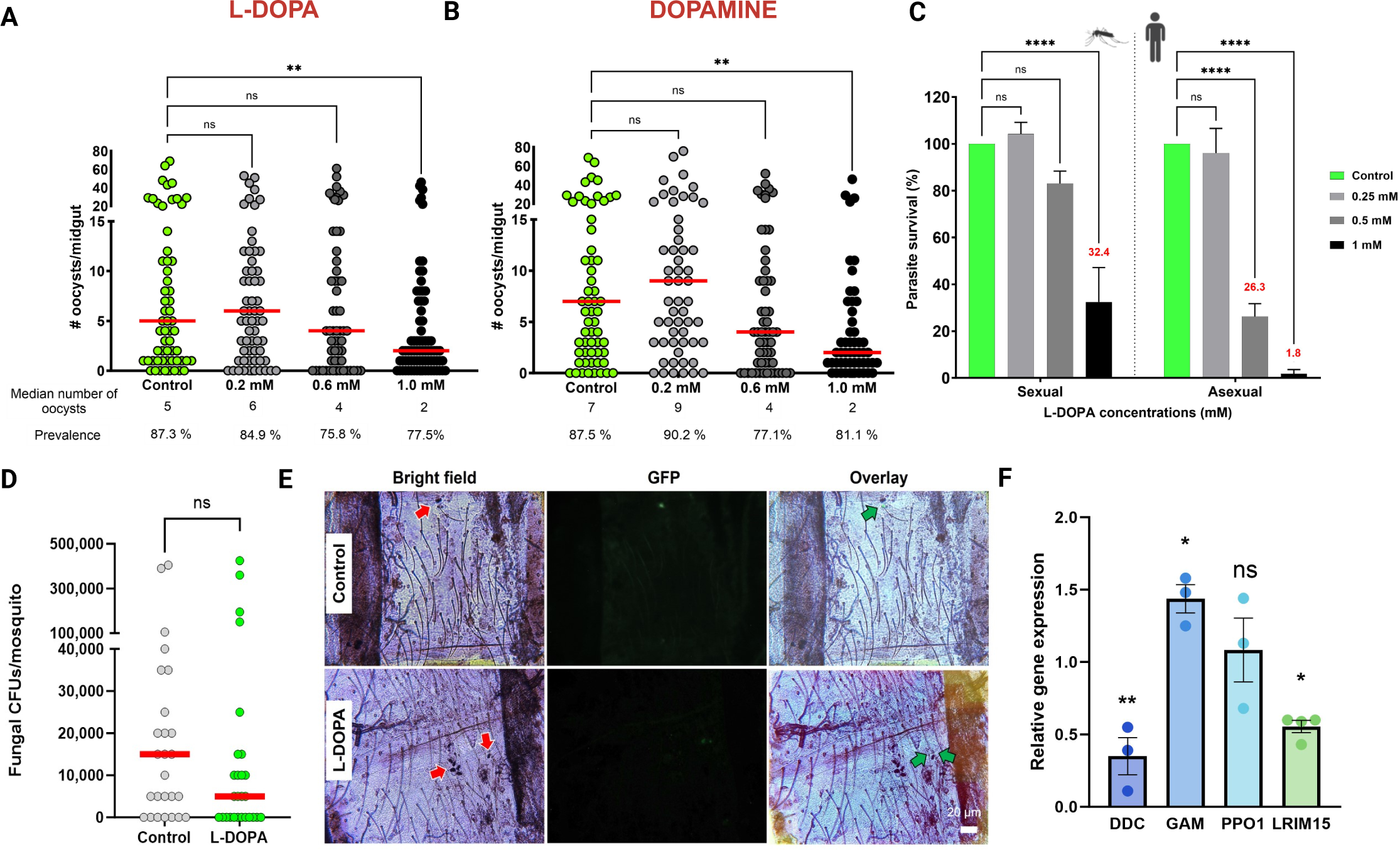
L-DOPA exhibits antiparasitic and antifungal activity. **a** *P. falciparum* infection intensity and prevalence in adult female mosquitoes fed on a sugar meal supplemented with a L-DOPA or dopamine concentrations. Each circle indicates the number of oocysts on an individual midgut; horizontal bars represent the median value of oocysts. Each experiment consisted of three biological replicates, and data were pooled to generate the graph. Statistical analysis of infection intensity was performed using the Kruskal-Wallis test with Dunn’s multiple comparisons post-test against the control; ns not significant, left panel \*\**p* = 0.0081, right panel \*\**p* = 0.0019. Statistical significance for prevalence of infection was assessed with a Fisher’s exact test, ns. **b** *In vitro* assessment of L-DOPA toxicity to sexual and asexual stages of *P. falciparu*m parasite. Use of L-DOPA concentrations ranging from 0.25 mM to 2 mM during *in vitro* parasite culture, appeared more toxic to asexual stages of the parasite than to sexual stages. Data from three biological replicates were pooled to generate the graph. Data was analyzed with ordinary two-way ANOVA with Dunnett’s multiple comparisons tests against individual controls; ns not significant; \*\*\*\**p*< 0.0001. **c** Fungal load in L-DOPA-fed female mosquitoes shows a reduction compared to control. Each circle corresponds to a whole mosquito homogenate; bars represent the mean value of fungal CFUs, and error bar corresponds to SD. Data from three biological replicates were pooled to generate the graph. **d** Cryptococcal yeasts are successfully melanized in the abdomens of *A. gambiae* female mosquitoes 24 h post-injection of *C. neoformans* H99-GFP strain (1 x 10^8^ yeasts/ml). Live yeast cells (green arrows), Dead yeast cells (white arrows). Representative images from abdomen sections of 10 female mosquitoes at two independent experiments. **e** qRT- PCR analysis of mRNA abundance in whole mosquitoes from key genes [dopa decarboxylase (DDC), gambicin (GAM), phenoloxidase 1 (PPO1), and leucine-rich immunomodulator 15 (LRIM15)] involved in the mosquito melanization pathway and *Plasmodium* parasite development. Data represent the mean ± SE of three or more independent replicates, analyzed using unpaired one-way ANOVA with Holm-Sidák’s multiple comparison test to determine gene expression relative to housekeeping S7 gene; ns not significant, \*\**p* = 0.0041, \**p* = 0.0235.

*A. gambiae* triggers a melanin-based immune response against fungi ^34,35^. To ascertain whether L-DOPA supplementation promoted fungal melanization, we challenged mosquitoes with an intrathoracic injection of *Cryptococcus neoformans* (H99-GFP strain), prior to L-DOPA exposure for 4 d. Fungal viability was assessed by measuring the number of fungal colony-forming units (CFUs) 24 h after injection and by monitoring H99-GFP fluorescence, a previously validated reporter strain for assessing fungal viability^36^. L-DOPA-fed mosquitoes showed reduced number of fungal CFUs in comparison to the control (Fig. 6c) and revealed efficiently melanized yeast cells (Fig. 6d). Furthermore, we found that despite downregulation of dopa decarboxylase (DDC), a key gene on the melanization pathway, other genes involved in *Plasmodium* parasite killing and survival —specifically the antimicrobial peptide gambicin (GAM)^37^ and the leucine-rich immunomodulator 15 (LRIM15)^38^— were significantly deregulated, affecting *Plasmodium* parasite development (Fig. 6e). Our analyses demonstrated that, despite significant downregulation of canonical melanization-related genes that was validated by qRT-PCR (Supplementary Fig. 2), L- DOPA-supplementation promoted both antiplasmodial and anti-fungal response in *A. gambiae* female mosquitoes.

### Dietary L-DOPA enhances mosquito cuticular melanization

The insect cuticle protects against physical trauma (rigid cuticle) and flexibility to support joint areas allowing mobility (soft cuticle) ^39^. Melanin is primarily thought to be involved in the formation of rigid cuticles, though its participation in the biogenesis of flexible cuticles is uncertain. While assessing daily mortality, we serendipitously noticed that the 1 mM L-DOPA-fed mosquitoes displayed a greater darkening of the cuticle pigmentation in comparison to sucrose-fed controls (Fig. 7a, upper panel). Thus, we explored the contribution of dietary L-DOPA to insect cuticular melanization by allowing *A. gambiae* female mosquitoes to feed with a sugar solution supplemented with L-DOPA (0.2 mM to 1 mM) for up to 14 d. At 8 and 14 d, we observed and measured a dose-dependent darkening of the mosquito cuticle; 1 mM L-DOPA-fed mosquitoes exhibited significantly darker pigmentation relative to sugar-fed control mosquitoes (Fig. 7a, lower panel). To further assess the mosquito cuticular ultrastructure and chemical nature of its pigment, we used transmission electron microscopy (TEM) to examine the cuticular architecture of legs from *A. gambiae* female mosquitoes after 8 d of exposure to the L-DOPA diet. Micrographs from female L-DOPA-fed mosquitoes revealed a distinct microstructure of the leg endocuticle layer in comparison to the sucrose-fed control (Fig. 7b, top panel). The mean thickness of the total cuticle was significantly greater (mean = 2,424.11 ± 467.97 nm) than that of the control (mean = 1,962.63 ± 302.91 nm) (Fig. 7b, bottom panel). In response to the L-DOPA diet, female mosquitoes showed cuticle enrichment promoted by an increase of both procuticle layers (endocuticle and exocuticle). In addition, we determined the accumulation of melanin in whole mosquitos using solid-state NMR with ^13^C-isotopically-labeled L-DOPA. Adult female *A. gambiae* mosquitoes were fed with 1 mM ring-^13^C-L-DOPA in sucrose for 8 d to augment the ssNMR signal intensity. To estimate the relative amounts of long-chain lipids, polysaccharides, and melanin in *A. gambiae* mosquitoes fed with or without labeled L-DOPA, we acquired quantitatively reliable ^13^C direct-polarization magic-angle spinning (DPMAS) NMR data. In comparison to the control, spectra from ring-^13^C-L-DOPA-fed female mosquitoes displayed increased signal intensities at 110-160 ppm which are characteristic of the indole-based melanin pigment (Fig. 7c, green arrows in left panel), suggesting that mosquitoes used ring-^13^C-L-DOPA to synthesize cuticular melanin. To determine whether mosquitoes broke down ring-^13^C-L-DOPA for nutritional purposes, we also acquired ^13^C cross-polarization magic-angle spinning (CPMAS) spectra that favored the most rigid constituents. This latter study corroborated the nearly exclusive use of ring-^13^C-L-DOPA for pigment production: signals corresponding to polysaccharides (∼55-105 ppm) as well as those corresponding to aliphatic carbons (10-40 ppm) were largely unchanged in integrated intensity upon feeding with ring-^13^C-L-DOPA (Fig. 7c, right panel). Interestingly, L-DOPA supplementation also increased the content of indole groups but reduced the cuticular thickness of legs in male *A. gambiae* mosquitoes (Supplementary Fig. 3). These mosquitoes also displayed a modest increase in their chitin content (0.64 ± 0.07 nmoles of glucosamine per mg of whole mosquito dry weight) in comparison to the sucrose-fed controls (0.46 ± 0.06 nmoles of glucosamine per mg of whole mosquito dry weight) (Fig. 7d). Furthermore, we tested whether the augmented indole-pigment content corresponded to melanin by performing electron paramagnetic resonance (EPR) measurements. EPR is a particularly effective methodology that detects a remarkable characteristic feature of this biopolymer, the presence of a population of stable organic free radicals ^40^. Analysis of the acid-resistant material isolated from L-DOPA-fed mosquitoes revealed a spin concentration that decreased to 0.81-fold of its value (14.55 vs 17.75, scaled by mass) compared with sugar-fed controls (Fig. 7e). Lastly, we demonstrated that melanin synthesis was not driven by DCE providing a L-DOPA diet to the *Aedes aegypti* yellow (DCE2, AAEL006830) mutant strain^41^, which is defective for eumelanin synthesis. This mutant mosquito strain also exhibited a spin concentration that diminished to 0.3-fold of its value (2.36 vs 7.60) compared with its sugar-fed control. The lower spin concentrations of L-DOPA supplemented samples in comparison to controls is suggestive of an enhanced radical scavenging activity— a key melanin property^42^. Notably, the high spin concentration exhibited by the *A. gambiae* L-DOPA supplemented samples compared with the *Aedes* L-DOPA-fed mutant mosquito strain (14.55 vs 2.36, respectively) could imply the isolation of different types of melanins (a mixture of pheomelanin/eumelanin vs eumelanin) with distinct antioxidant properties. These results agree with our fungal eumelanin sample, used a control, which displayed a spin concentration of 4.26. Further studies will be required to gain a deeper understanding of this system.

**Fig 7.**
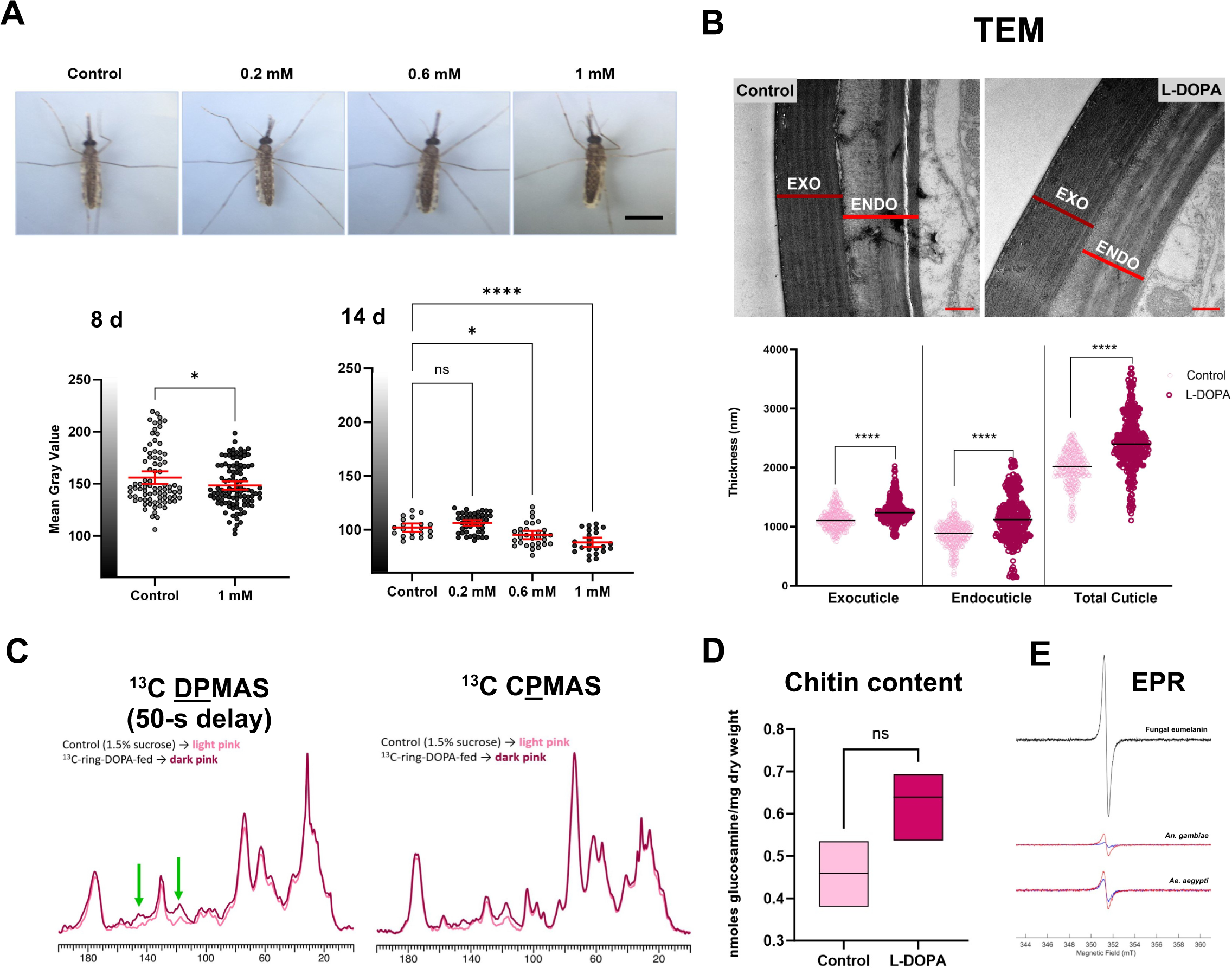
Oral L-DOPA boosts mosquito cuticular melanization via a non-canonical pathway. **a** Dietary L-DOPA enhances cuticle pigmentation of *A. gambiae* female mosquitoes in a dose- dependent manner. *Upper panel*, Representative images of *A. gambiae* female mosquitoes fed with a sugar meal supplemented with different concentrations of L-DOPA for up to 14 d. Scale bar, 1 mm; *Lower panel*, Quantification of cuticle darkening at 8 and 14 d. Each dot corresponds to an individual mosquito. Mean with 95% CI from 2-3 biological replicates are shown in graphs. Statistical significance was determined by Left panel, unpaired two-tailed *t*-test, \**p* = 0.0342; and Right panel, ordinary one-way analysis of variance (ANOVA) with Dunnett’s multiple comparison test against control, ns not significant, \**p* = 0.0365, \*\*\*\**p* < 0.0001). **b** *Top panel,* TEM micrographs exhibiting distinct electron-dense bands within the endocuticle of L-DOPA-fed mosquitoes that are not seen in the control. Scale bar, 500 nm. *Lower panel,* Measurements of cuticular thickness from legs of female mosquitoes fed with L-DOPA supplemented sugar meals and control show increased values for exo and endocuticle. Two-way ANOVA with Tukey’s multiple comparisons test; \*\*\*\**p*<0.0001. Data represent measurements from 10 femurs per condition with 3-5 representative images analyzed per female. A minimum of 3 total cuticle and endocuticle measurements were taken per image. Means with SD are shown on the graph. **c** L-DOPA supplementation augments content of indole groups. *Left panel*, DPMAS ^13^C ssNMR spectra of ^13^C-L-DOPA fed mosquitoes display pigment peaks (green arrows) with greater relative intensities than their respective controls. Spectra were normalized to the tallest peak (lipid peak at ∼30 ppm); Right panel, CPMAS ^13^C ssNMR spectra of ^13^C-L-DOPA fed mosquitoes corroborate the assertion that ^13^C-L-DOPA has been nearly-exclusively used to make melanin instead of being broken down and used as a nutrient source. Spectra were normalized to the tallest peak (polysaccharide peak at ∼72 ppm). Representative data from 2 biological replicates. **d** Quantification of chitin content in whole *A. gambiae* female mosquitoes demonstrates no significant differences between sugar-fed (control) and L-DOPA-fed mosquitoes. Data correspond to 3 biological replicates. Mean with SD. Two-tailed unpaired, nonparametric Mann-Whitney test; ns not significant. **e** EPR spectroscopy analysis of acid-resistant material isolated from *An. gambiae* and *Ae. aegypti* yellow mutant strain L-DOPA-fed (red spectra) and sugar-fed (blue spectra). Representative data from 2 biological replicates. Fungal eumelanin was used as positive control (black spectrum).

In sum, oral L-DOPA promoted mosquito cuticle melanization via a non-canonical pathway independent of L-DOPA conversion into dopachrome, impacting the thickness of both endocuticle and exocuticle layers.

### Enhanced cuticle melanization fails to promote insecticide resistance yet raises mosquito body temperature and reduces lifespan

Previous studies in *Anopheles* mosquitoes have shown that changes in the cuticle structure are associated with insecticide-resistance ^43,44^. Thus, we set out to determine whether enhanced cuticular melanization impacted female *A. gambiae* resistance to pyrethroids, the only WHO-approved insecticide used on long-lasting insecticide-treated nets (LLINs). Using WHO-approved papers impregnated with 0.05% deltamethrin for 1 h exposure, L-DOPA-fed *A. gambiae* female mosquitoes demonstrated susceptibility to pyrethroids (98.7± 3.3% mortality rate in 24 h). Despite their highly melanized and thicker legs, L-DOPA-fed *A. gambiae* female mosquitoes were susceptible to 0.05% deltamethrin.

Dark pigmentation is associated with enhanced ability to capture heat from electromagnetic radiation on animals, plants, and fungi ^45^. In insects, melanization (melanism) is associated with negative or positive consequences in thermal adaptation, mating, cuticle strength, etc. that impact their fitness and ecology ^46–48^. To probe mosquitoes’ thermal response to visible electromagnetic radiation, we used a thermal camera to measure the maximum average body temperature of L-DOPA-fed individual mosquitoes after being exposed to a white light mimicking sunlight. Female *A. gambiae* mosquitoes fed with L-DOPA for 8 d maintained an apparent average body temperature that was 0.5 – 2.0 °C higher that sugar-fed control mosquitoes, following 1 min of irradiation (Fig. 8a, Supplementary Fig. 4). Notably, 1 mM L-DOPA-fed mosquitoes were significantly warmer than control. These mosquitoes exhibited the highest temperature, both pre- and post-irradiation (Fig. 8b, Supplementary Fig. 4). Because trade-offs between heat tolerance and life-history are observed in insects ^49^, we further investigated whether longevity of L-DOPA-fed mosquitoes differed from those of controls. We measured the longevity of female mosquitoes when maintained on the L-DOPA diet (Fig. 8c). The lifespan of female mosquitoes was significantly decreased when maintained on a sucrose meal supplemented with 1 mM L-DOPA (Wilcoxon test, Chi-square, 13.74, *p* = 0.0002). These mosquitoes exhibited no apparent changes in reproductive fitness: number of eggs laid (Fig. 8d) and hatching rate (Fig. 8e).

**Fig. 8.**
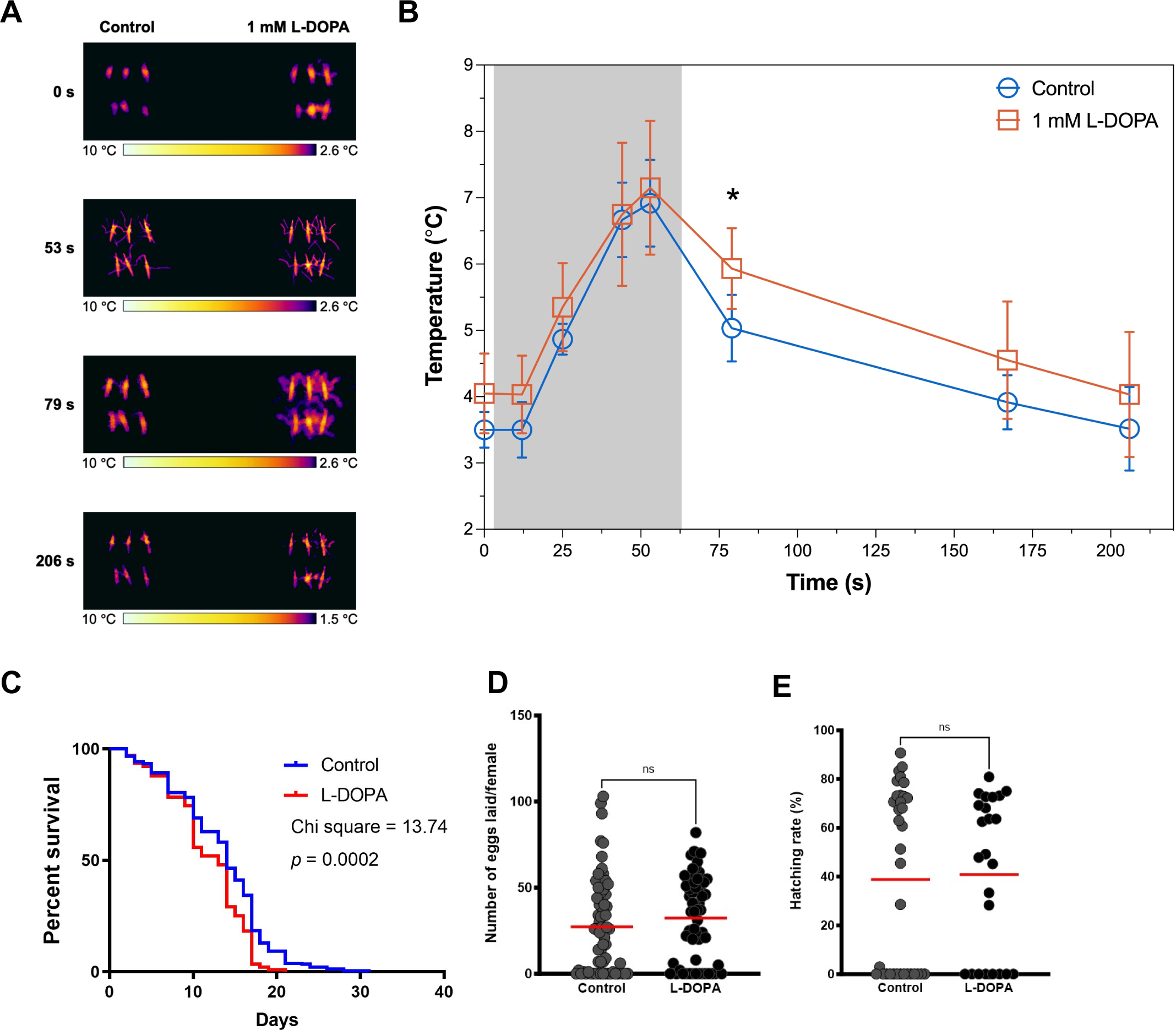
Dietary L-DOPA darkening of cuticle increases heat-exposed mosquito temperatures following light exposure and decreases lifespan. A) *A. gambiae* female mosquitoes fed with a sugar meal supplemented with different concentrations of L-DOPA (0.2 mM to 1 mM) for 8 d displayed a dose-dependent ability to absorb heat. **B)** Exposure to a direct light source during 1 min (shadowed area) demonstrated that 1mM L-DOPA- fed mosquitoes (red squares) exhibited a warmer body temperature than sugar-fed control (blue circles). Post- irradiation when light is turned off (unshadow area), L-DOPA-fed mosquitoes (red squares) were distinctly hotter than sugar-fed controls (blue circles), starting to cool down but kept warmer than control The thermograph was obtained using the Teledyne FLIR E95 camera, and the temperature of each mosquito was extracted using the FLIR software, which provided the maximum, minimum, and average temperatures within a box selection for each mosquito. The maximum temperature for each mosquito was used to generate the graph, which represents the average (+/- Standard Deviation) of the six maximum temperatures corresponding to the two experimental groups (control sugar-fed and 1 mM L-DOPA-fed). Mann-Whitney test, *p* = 0.0173. Mann-Whitney test, \**p* = 0.0173. **C)** Lifespan of mosquitoes maintained on 1.5% sugar with 1 mM L-DOPA was diminished in comparison to sugar-only fed mosquitoes. Representative data from 3 biological replicates. **D)** Number of eggs laid by female mosquitoes after feeding with a sugar meal supplemented with 1 mM L-DOPA during 4 d before a blood meal on mice. Mann-Whitney test, NS, not significant, *p* = 0.1987. Red lines denote mean. **E)** Hatching rate at 27 °C indicating average percentage of eggs giving rise to 1st and 2nd instar larvae. Mann-Whitney test, NS, not significant. *p* = 0.9390. Red lines denote mean.

### The mosquito microbiome includes bacteria that produce dopamine

Unlike mammalian tyrosinases, which utilize L-DOPA as a melanin precursor ^50^, insect phenoloxidases preferentially use dopamine over L-DOPA ^51^. This led us to investigate how dietary L-DOPA was metabolized in female mosquitoes and whether the mosquito microbiome harbors dopamine-synthesizing bacteria. Contrary to findings in *Aedes albopictus* ^52^, we observed the intriguing trend whereby as L-DOPA levels increased in the sugar meal, dopamine levels in mosquitoes decreased (Fig. 9a). Leveraging our experience with studying fungal melanization in *Cryptococcus* spp., which can be triggered by dopamine synthesized by bacteria ^53^, we explored whether gut microbiota isolated from field-caught mosquitoes have the potential to serve as an endogenous source of dopamine. *Cryptococcus* spp., known to be members of the mosquito gut mycobiome ^54,55^, depend exclusively on exogenous precursors for melanin synthesis (e.g., L-DOPA, dopamine, epinephrine, and norepinephrine) ^56^. We selected thirteen bacterial strains that had been previously isolated from the midgut of field-caught *Anopheles* ^16^ and *Aedes* ^57^ mosquitoes plus a strain isolated from the soil^58,59^ (Supplementary Table 1) and conducted a plate confrontational assay whereby bacteria that produce dopamine trigger melanization in *Cryptococcus neoformans* ^53^. After eye inspection of the plates, we noticed that five bacterial isolates consistently induced melanization (brown pigmentation) in both of our positive control strains (*C. gattii* WM179 and *C. neoformans* H99), at both room temperature and 30°C. As expected, no pigmentation was observed in the *C. neoformans* LAC1 mutant strain (negative control), in which melanin synthesis is compromised by the deletion of its laccase gene (Fig. 9b). The melanin-inducing bacteria were identified as: *Comamonas* sp., *Enterobacter hormaechei*, *Aeromonas hydrophila*, *Serratia marcescens*, and *Chromobacterium subtsugae ΔvioS* (*ΔvioS*). To verify the chemical nature of the soluble compound released by mosquito-derived bacteria that promoted cryptococcal melanization, we used a Fast Protein Liquid Chromatography (FPLC) protocol for recovering dopamine from bacterial supernatants grown in Luria-Bertani (LB) media for up to 96 h at 30°C. The chromatograms displayed a peak between 4.39 and 5.66 min matching the position of the dopamine spike peak. We observed that peaks varied depending on the bacterial species and tended to be larger as the incubation time increased (Supplementary Fig. 5). Therefore, we collected peaks at 96 h of culture incubation, removed cells by quick centrifugation, filter-sterilized the supernatants, acidified them to pH 4.0 with 1 M HCl to maintain dopamine stability, and kept them on ice, and away from light. Samples were further identified as dopamine and quantified by Electrospray Ionization (ESI) Mass Spectrometry (MS). Dopamine production was verified from four strains at concentrations ranging from 7 ng/ml to 67.5 ng/ml. These were the dopaminergic bacteria identified (concentration of dopamine detected): *Aeromonas hydrophila* (7.10 ± 0.4 ng/ml), *Comamonas sp.* (7.00 ± 0.1 ng/ml) and *Enterobacter hormaechei* (67.5 ± 2.4 ng/ml). No dopamine was quantified for *Serratia marcescens* under the tested conditions. The limit of quantification of the ESI MS was 2.5 ng/ml.

**Fig. 9.**
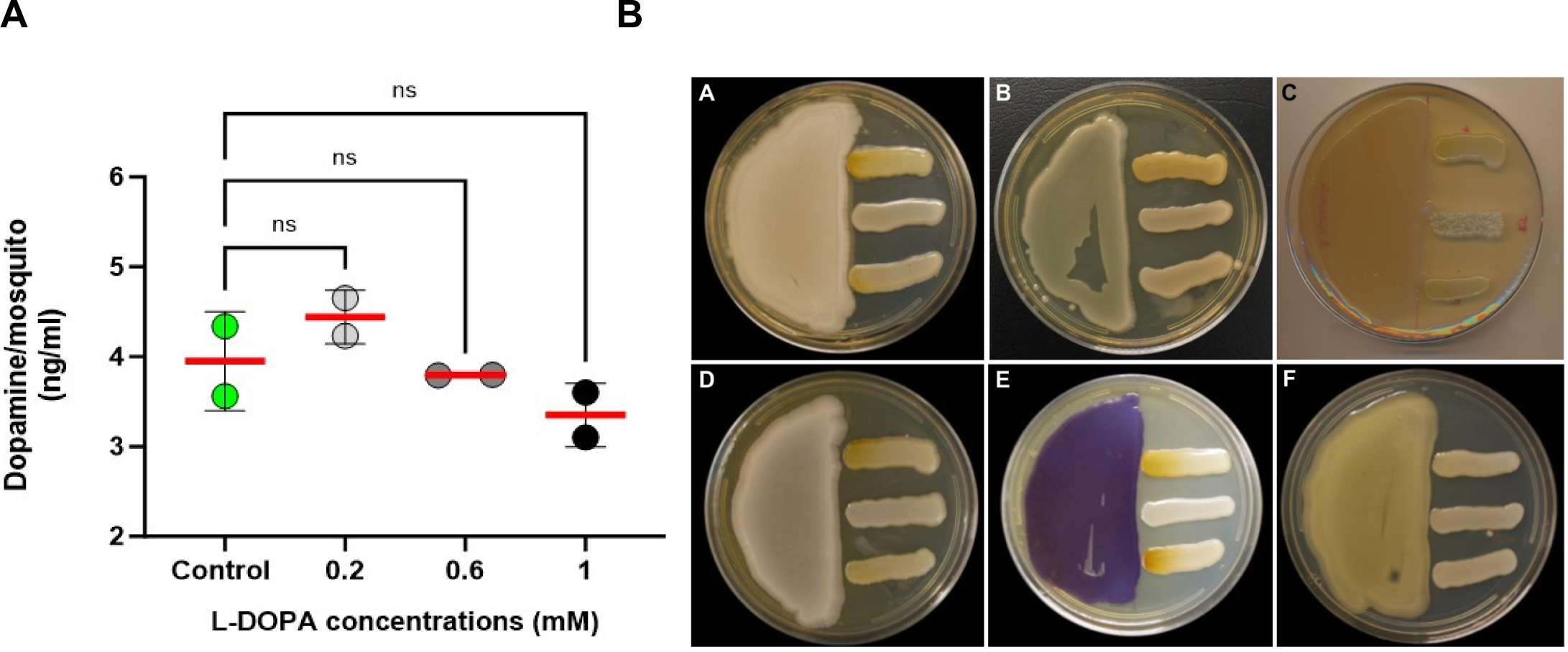
D**i**etary **L-DOPA metabolism and probing for dopaminergic bacteria amongst gut microbiota from field-caught mosquitoes. A)** Quantification of dopamine by HPLC from whole *Anopheles* mosquitoes post 4 d of dietary L-DOPA suggests tight regulation of L-DOPA conversion into dopamine. Data correspond to 2 independent biological replicates from 5-pooled mosquitoes per condition. A 2-way analysis of variance (ANOVA) with Tukey’s multiple comparisons test showed no significant differences among and between groups. **B)** Plate confrontational assays showed induced fungal melanization due to melanin precursors released by bacterial isolates. A) *Comamonas* sp.; B) *Enterococcus hormaechei*; C) *Aeromonas hydrophila*; D) *Serratia marcescens* E) *Chromobacterium subtsugae ΔvioS* F) *Enterococcus* sp. *Cryptococcus* strains (top: *C. gattii* WM179; middle: *C. neoformans* ΔLAC1; bottom: *C. neoformans* H99) were streaked perpendicular to bacterial isolates in SM plates. The plates were incubated at 30°C for 5-7 days. Images are representative of 3 biological replicates.

## Discussion

In insects, melanin is synthesized both in the exoskeleton (cuticle) and inside the body, including by the immune system, which uses melanization as an antimicrobial mechanism ^60–62^. The cuticle is crucial for insect survival, making it a primary target for pest and disease vector control. Its mechanical properties (i.e., rigidity and flexibility) are related to various chitins, proteins, and the type and abundance of sclerotization or protein crosslinking occurring within ^63^. This study demonstrates that *Anopheles* mosquitoes can use dietary L-DOPA to enhance cuticle melanization, leading to increased body pigmentation, cuticular thickness in the legs, overall indole and chitin content, and a key EPR signal (Fig. 7). Given the abundance of L-DOPA in the environment and the potential microbiome sources of dopamine, our results have important implications for mosquito biology and potential control strategies.

Cuticular melanogenesis is closely related to cuticular hardening or sclerotization. Both are oxidative processes occurring in the same tissues and use L-DOPA or dopamine as precursors; however, the participation of laccase 2 (AGAP008731) and dopamine derivatives such as *N*- acetyl-dopamine (NADA) and *N*-β-alanyldopamine (NBAD) in mediating covalent crosslinks between cuticular proteins and/or between cuticular proteins and chitin fibers is exclusive to the sclerotization process ^17,51^. Our data demonstrate that mosquitoes can metabolize L-DOPA via a non-canonical pathway using 3,4-dihydroxylphenylacetaldehyde (DOPAL) synthase (Fig. 10a). As a detoxification mechanism, L-DOPA is directly converted into DOPAL and hydrogen peroxide. The resulting hydrogen peroxide further oxidizes the DOPAL catechol ring, leading to the formation of a melanized and flexible cuticle crosslinked with cuticular proteins, while bypassing the conversion of L-DOPA to dopamine ^63,64^. The buildup of DOPAL promotes oligomerization and the formation of quinone adduct (“quinonization”), which inhibits its own metabolism and the activity of other enzymes involved in the melanin biosynthesis ^65^. Therefore, this study supports that L-DOPA-induced downregulation of key melanization pathway genes (e.g., *DDC* and *DCE*) occurs at transcriptional level leading to a reduction of their protein translation/production (Fig. 10b). Further studies are required to demonstrate DOPAL buildup and protein modification.

**Fig. 10.**
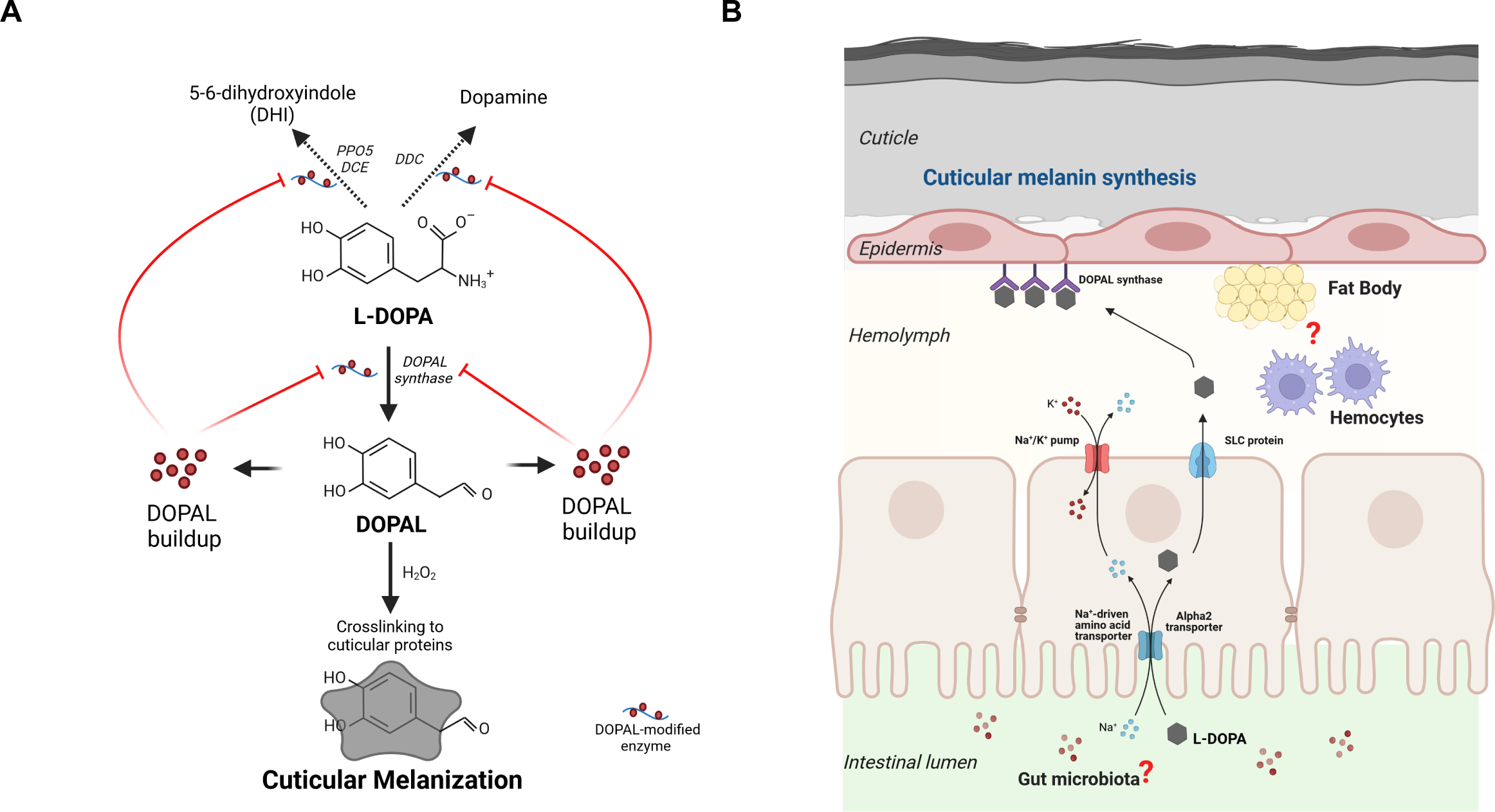
A**l**ternative **pathway for L-DOPA incorporation to the melanin synthesis in mosquitoes. A)** L-DOPA serves as a substrate for cuticular melanization via DOPAL synthase as a detoxification mechanism given a cellular redox environment that promotes L-DOPA antioxidant properties (neutral or acid pH) ^66^, therefore preventing its conversion to dopamine via a dopa decarboxylase. L-DOPA conversion into DOPAL leads into DOPAL buildup that promotes oligomerization and quinone adduct formation (“quinonization”) inhibiting its own metabolism and the activity of other enzymes involved in the melanin biosynthesis such as dopa decarboxylase (DDC), phenoloxidase 5 (PPO5), and dopachrome conversion enzyme (DCE)^65^. **B)** Model for dietary L-DOPA-mediated cuticular melanization proposes that *Anopheles gambiae* female mosquitoes use L-DOPA supplement in the sugar meal as a substrate for synthesizing cuticular melanin. L-DOPA is internalized into the midgut epithelium and exported to the hemolymph by sharing the same transporters with amino acids such as Na^+^-driven amino acid transporter, alpha2 transporter, and/or SLC protein. Within regions of flexible cuticle, in the presence of DOPAL synthase, L-DOPA is turned into DOPAL and H2O2 promoting oxidation of DOPAL catechol ring that results in the formation of a melanized and protective flexible cuticle via crosslinking to cuticular proteins^63,64^. The contribution of gut microbiota as well as from hemocytes as key regulators of mosquito vectorial capacity and immunity, post a L-DOPA diet is under investigation. Created in BioRender. Camacho, E. (2025) https://BioRender.com/d90z642

Dietary L-DOPA affects multiple mosquito physiological pathways in a dose-dependent manner (Figs. 3, 6, 7, 8, 9). The strong allelopathic activity of L-DOPA may explain our observations ^27^, Depending on the cellular redox potential, L-DOPA behaves as a prooxidant (at alkaline pH) promoting oxidative damage and covalent modification of proteins, or as an antioxidant (at acid and neutral pH) scavenging initiating radicals by electrostatic interactions and chelating metal ions to prevent their participation in radical formation ^66^. Our analyses demonstrate that L-DOPA-supplemented sucrose meals induced a biphasic dose-response (Fig. 3b), a phenomenon known as hormesis. Hormesis occurs in response to moderate environmental or self-imposed challenges, which causes a system to improve its functionality and/or tolerance when faced with more severe challenges ^67,68^. This phenomenon is widespread in nature and has been reported for various molecules, including nitric oxide, prostaglandins, dopamine, and other molecules^69^. Hormetic-like responses involve the simultaneous stimulation of many independent cellular functions/endpoints and define the boundaries of biological plasticity ^68^.

Despite marked downregulation of immune-related genes (Fig. 5a), oral L-DOPA promoted antimicrobial activity in *A. gambiae* female mosquitoes (Fig. 6). Our *in vitro* analyses show that L-DOPA concentrations above 0.5 mM impair *P. falciparum* development at both sexual and asexual stages. This aligns with a previous finding that L-DOPA induces apoptosis in *P. berghei* ookinetes under similar conditions ^70^. Conversely, given the particularity of our L-DOPA diet (an amino acid analog within a low sucrose concentration) and the complexity of the mosquito digestion system (sugar and protein meals have distinct compartmentalization) ^71^, the enhanced resistance to parasite burden exhibited by L-DOPA-fed mosquitoes might reflect multifactorial mechanisms. At least two possible scenarios, non-mutually exclusive, could explain the results. First, L-DOPA-mediated acidification of midgut pH could block parasite development at the midgut lumen thus reducing midgut invasion ^72^; mean pH measurements of 1 mM L-DOPA-sugar solution dropped from 8.3 to 6.7 within 72 h (Supplementary Fig. 6). Second, the restrictive sugar diet (1.5% sucrose) may trigger a metabolic switch that functionally primes the mosquito to resist pathogen invasion ^73^. In line with the latter scenario, the upregulation of the Dally protein (AGAP005799), phosphoinositide-3-kinase (PI3K) (AGAP005583), and juvenile hormone (JH) (AGAP008685) could reflect an integrative response from the insulin/TOR signal transduction pathway, sensing not only nutritional information but also other environmental cues such as temperature and foreign organisms to fine-tune allocation of resources and promote fitness ^74–77^.

In most insects, genes encoding enzymes involved in melanization and sclerotization are referred to as pleotropic ^78–80^, supporting the hypothesis that pleiotropy (i.e., expression of multiple traits by a single gene) drives the coevolution of correlated traits ^81^. Such is the case for cuticle pigmentation, which was significantly darkened in 1 mM L-DOPA-fed mosquitoes (Fig. 7). Its association with increased thickness in female mosquitoes immediately raises a concern for insecticide resistance given that several studies in disease vectors have reported this linkage ^43,82–84^. Other mechanisms extensively shown to be involved in resistance to insecticides are alteration of target molecule, reduced cuticle permeability, and metabolic clearance via overexpression of detoxification protein families such as cytochrome P450 (CYPs), glutathione S transferases (GSTs), and esterases ^85^. In agreement with *A. gambiae* susceptibility to deltamethrin upon L-DOPA feeding, these mosquitoes exhibit a drastic downregulation of GSTs, carboxylesterases, cuticular proteins, and CYPs including CYP4C27 (AGAP009246), which were previously associated with insecticide resistance in *A. gambiae* ^86^ (Fig. 5b, Supplementary Fig. 2). It is worth mentioning that the slight increase of chitin content measured in whole L-DOPA fed mosquitoes corresponds to a mixture of chitin and chitosan, given that the samples were deacetylated before performing the colorimetric reaction. Regardless of the chemical processing, based on our previous work with melanized cryptococcal cell walls ^42,87^, as well as the observed downregulation of chitin synthase 1 (CHS1A,B) (AGAP001748), we propose that dietary L-DOPA induces an increase of chitosan within the mosquitoes legs. This, in turn, promotes melanin deposition and thus translates into augmented cuticular pigmentation. As predicted by the thermal melanism hypothesis ^88^, increased mosquito pigmentation was associated with augmented heat absorption for the 1 mM L-DOPA fed mosquitoes (Figs. 8a-b) supporting an adaptative trait also described in insects from various orders ^89–91^. *Anopheles* mosquitoes are poikilotherms, so ambient temperature strongly influences developmental processes, behaviors, and even survival ^92^. These effects in thermoregulation could have an ecological impact; indeed, darker mosquitoes displayed a reduced longevity (Fig. 8c). This negative effect could be attributable to low tolerance to heat associated with starvation^49^ given that our polyphenol-rich diet (e.g., dietary L-DOPA) with low sucrose concentration mimics calorie restriction ^93^. Another possible explanation for the decreased life span is that dietary L-DOPA might be affecting *A. gambiae* cuticular hydrocarbon composition (CHC), therefore increasing susceptibility to desiccation. This reasoning is based on our RNA-seq data from L-DOPA-fed *A. gambiae* female mosquitoes, which reveals a strong downregulation of genes involved in CHC synthesis, including three genes specifically enriched in *A. gambiae* oenocytes (immune cells associated to melanin production) (Figure 5B). One of these genes, AGAP001473 [3.05-fold change, *p* = 0.0004 compared to sugar-fed controls, validated by qRT-PCR (Supplementary Fig. 2), encodes a single propionyl-CoA synthetase responsible for synthesizing methyl-branched hydrocarbons (mbCHCs) ^94^. These mbCHCs have melting points (temperature at which these hydrocarbons transition from a solid to a liquid state) above ambient temperature, meaning they likely play a role in both reducing water loss and mediating chemical communication ^95^. We hypothesize that in response to dietary L-DOPA, *A. gambiae* female mosquitoes exhibit a distinct CHCs profile, skewed toward short-chain (<25C) and non-branched CHCs, leading to increased susceptibility to desiccation. Further experimental testing will be required to test this hypothesis.

Concomitant with dietary L-DOPA, our evidence that gut microbiota colonizing adult *Aedes* and *Anopheles* mosquitoes can synthesize dopamine (Fig. 9b) strongly suggests that the mosquito gut microbiome could also play a role in the effects reported here. Indeed, certain *Enterococcus* and *Bifidobacterium* species identified in in the gut of Parkinson’s Disease patients can metabolize L-DOPA into dopamine or other metabolites like 3,4-dihydroxyphenyl pyruvic acid (DHPAA) ^96,97^. The mosquito microbiome is a critical modulator of vector competence and vectorial capacity, also influenced by the mosquito diet ^16,98,99^. The interactions between fungal and bacterial microbes shape fungal microbiome communities within the mosquito gut, which are dominated by Ascomycota and Basidiomycota. Among these are *Candida* and *Cryptococcus* spp.^54,55^ which can use catechols (L-DOPA and dopamine) as substrates for melanin synthesis ^56^. Therefore, as previously reported by other authors ^100–102^, midgut microbiota could also contribute to mosquito homeostasis by detoxifying toxic compounds that modulate symbiotic relationships between the gut microbiome and their host. In this line of thinking, the L-DOPA-modulated microbiome can also promote a certain pH at the midgut, thus favoring either prooxidant or antioxidant properties of L-DOPA ^72^. Interestingly, recent research in *Aedes aegypti* demonstrated that higher levels of juvenile hormone (JH) in mated and virgin-JH treated in comparison to virgin mosquitoes promoted establishment of the midgut microbiota, midgut enlargement and allocation of energetic resources to enhance fecundity while inducing some immunosuppression. L-DOPA-fed female mosquitoes had no defects in reproduction (Figs. 8d-e). Recent evidence in the invertebrate model *Daphnia magna* showed that its gut microbiome is responsible for the synthesis of L-DOPA, which may serve as a signaling molecule modulating host-microbiome interactions relevant for key life traits ^103^. The existence of a gut-to-brain signaling axis modulated by 5-hydroxytryptamine innervations in the midgut tissue of *Anopheles stephensi* mosquitoes was also recently suggested ^104^. In addition, the contribution from hemocytes as key regulators of mosquito immunity cannot be ruled out. In *Drosophila*, hematopoietic progenitors can directly sense and respond to systemic (sugar/insulin) and nutritional (amino acids) signals via coordinated communication between organs ^105^.

Furthermore, insect herbivores can use plant-synthesized compounds by ingesting and storing them in their hemolymph, integument (cuticle), and other compartments such as cuticular cavities, in subcuticular compartments, fat body, or in exocrine defense glands ^106^. This process, known as chemical sequestration, requires the active transport of the compound from the gut lumen across the midgut epithelium and to other compartments via ATP-binding cassette (ABC) and solute carrier (SLC) transporters ^107^. Unlike metabolic resistance, which involves alteration and detoxification of the compound or plant toxin, sequestered compounds must preserve their pharmacological activity, allowing insects to benefit from them. For instance, sequestered aromatic compounds have been linked to defense, reproduction, or exoskeleton stabilization in various insect orders, including Diptera^108^. Our RNA-seq data from midguts of L-DOPA-fed mosquitoes reveals upregulation of AGAP006221, a gene coding for an aldehyde oxidase, as well as two transport-related genes—sodium-dependent nutrient amino acid transporter 5 (AGAP010857) and nicotinic acetylcholine receptor subunit alpha 2 (AGAP002972).

AGAP006221 is highly upregulated in *A. stephensi* at early stages (up to 24 h) post permethrin exposure, thereby playing a critical role in its detoxification process^109^. Interestingly, in response to the insecticide-mediated stress, these authors found a broad downregulation of CYPs, GSTs, and carboxylesterases suggestive of reallocation of energetic resources during insecticide stress, as we also demonstrated in *A. gambiae* post dietary L-DOPA exposure. Given the pleotropic effects of dietary L-DOPA in *Anopheles* mosquitoes’ biology, supported by its strong allelochemical activity and potential direct role as a neurotransmitter mediating host-gut microbiome interactions ^103^, it was out of the scope of this work to elucidate the mechanism(s) responsible for the reduction of dopamine levels in whole mosquitoes as L-DOPA concentrations in the sugar increased (Fig. 9a). However, in agreement with our findings, L-DOPA sequestration with the same dopamine trend has also been observed in the pea aphid (*A. pisum*) when feeding on an L-DOPA-rich plant, the broad bean (*V. faba*). This phenomenon enhances wound healing and confers UVA-radiation protection to the pea aphid ^18^. A potential explanation for this could involve a gut symbiont (bacteria, fungi, or a combination of both) that modulates a mechanism to maintain mosquito homeostasis. This mechanism might promote L-DOPA detoxification while avoiding its conversion into dopamine, thereby regulating critical downstream processes in insects, such as sclerotization, melanization, and the synthesis of neurotransmitters (e.g., tyramine and octopamine)^17,110^. Additionally, this scenario might align with the potential modulation of gut microbiome composition, favoring the establishment of organisms that create a cellular redox microenvironment, which enhances L-DOPA antioxidant properties. Further research is needed to validate L-DOPA sequestration in *A. gambiae* and to assess its impact on reproduction, exoskeleton stabilization, and protective role against environmental stressors such as temperature, relative humidity, and UVA damage.

Our findings could be incorporated into mosquito control strategies. Despite strenuous measures to mitigate vector-borne diseases, these still account for more than 700,000 deaths annually, of which over 50% are caused by malaria - 96% of them in Africa ^111^. As proposed by other groups, the use of genetically modified mosquitoes to control malaria offers great hope but also comes with concerns and controversy ^112,113^. To effectively address these global risks and safeguard the biosphere on which all species depend, it is imperative to develop novel, safe, and biodegradable insecticides. Solutions derived from natural plant products hold promise in mitigating the impact of chemical pesticides on the environment. L-DOPA, the plant phenolic precursor of dopamine, is a neurotransmitter involved in several complex behaviors and processes in invertebrates, including locomotion, learning, courtship, development, and melanin synthesis ^51,114,115^. Native plants from Africa such as banana ^26^ (*Musa balbisiana* Colla) and Fava beans (*Vicia faba)* ^21,25^, contains up to 5% of L-DOPA in their leaves, flowers, and fruit parts. This metabolite is also a non-protein amino acid usually shown to be toxic to many insects ^66,116^ that could drive trade-offs between enhancing tolerance and life-history traits ^18,19,114^. By fostering the development and implementation of eco-friendly alternatives such as L-DOPA supplemented sugar baits designed to specifically lure defined mosquito species and using natural body odor chemoattracts ^117^, it may be possible to develop new tools for control of mosquito populations. Moreover, L-DOPA supplementation of mosquito-infested waters could enhance their immunity to reduce parasite burden and shorten lifespan, thus interfering with lifecycle and reducing the opportunity for parasite transmission.

In summary, feeding of a sugar meal supplemented with L-DOPA to adult *A. gambiae* mosquitoes augmented cuticular melanization and reduced parasite burden and lifespan. In addition to direct L-DOPA effects, increased melanization has the potential to affect insect thermoregulation. Consequently, L-DOPA supplementation has the potential for development as a novel, effective, and environmentally safe strategy to eliminate and/or reduce malaria transmission, which merits additional investigation including field studies of its usefulness and feasibility.

## Methods

### Ethical statement

The Johns Hopkins University Animal Care and Use Committee has approved this protocol, with permit number MO18H82. Commercial anonymous human blood was used for parasite cultures and mosquito feeding, and informed consent was therefore not applicable. The Johns Hopkins School of Public Health Ethics Committee has approved this protocol. Mice for mosquito blood feeding were housed in environmentally controlled rooms at 72°F and 42% humidity, with a 14.5- h light/9.5-h dark cycle.

### Mosquito rearing

Mosquitoes were obtained from colonies of *A. gambiae* (Keele strain) maintained in the insectary of the Johns Hopkins Malaria Research Institute (JHMRI) on a 10% sucrose solution with 12-h light/dark cycles at 27°C and 80% humidity under normal, non-sterile conditions. For most experiments in this study, except for the oral toxicity analyses, newly emerged control and treated mosquito groups were provided with 1.5% sucrose solutions in the absence or presence of the melanin precursor supplement.

### Plasmodium falciparum culture

A. *P. falciparum* NF54 asexual blood stage cultures were maintained *in vitro* according to the method described earlier by Trager and Jenson (PMID 351412). Briefly, parasites were cultured in O+ erythrocytes at a 4% hematocrit in RPMI 1640 (Corning) supplemented with 2.1mM L-glutamine, 25mM HEPES, 0.72 mM hypoxanthine (Sigma), 0.21% w/v sodium bicarbonate (Sigma), and 10% v/v heat-inactivated human serum (Interstate blood bank). Cultures were maintained at 37°C in a glass candle jar. Gametocyte cultures were initiated at 0.5% asexual blood stage parasitemia and at 4% hematocrit and maintained by changing the media daily for 17 d without the addition of fresh blood to promote gametocytogenesis as described earlier ^118^. Erythrocytes used for the study were obtained from healthy donors under a Johns Hopkins Institutional Review Board-approved protocol and were provided without identifiers to the laboratories.

### L-DOPA supplementation into sugar meal

L-DOPA (3,4-Dihydroxy-L-phenylalanine) (Cat N° D9628) was obtained from Sigma-Aldrich. Sterile sucrose solutions at 3% were diluted with an equal volume of sterile L-DOPA solutions and prepared from a 5 mM stock in distilled water to obtain sugar solutions supplemented with L-DOPA at concentrations ranging from 0.2 to 10 mM (0.4 to 20% w/w) L-DOPA. As control, 1.5% sucrose solutions were prepared. This low sucrose concentration setup stemmed from an unintentionally overdilution of sugar meals that revealed interesting and reproducible results, which we decided to analyze.

### Survival assay

Three- to four-day-old adult female *A. gambiae* were allowed to feed on sugar solutions prepared as described above. Survival of mosquitoes was monitored for 14 d, and dead mosquitoes were removed daily. Solutions containing 1.5% sucrose (end concentration) with an equal ratio of L-DOPA were freshly prepared and provided every 2-3 d. To facilitate handling of paper cups with mosquitoes (a group of 7, including sugar-only control), all cups were contained inside an airtight plastic container and then incubated at 27°C inside a reach-in incubator.

### RNA extraction and transcriptional profiling

Newly emerged adult female mosquitoes were allowed to feed with sugar solutions in the presence and absence of 1 mM L-DOPA for 4 d. After this, mosquitoes were cold anesthetized to remove their head using a tweezer over a glass plate on ice. Four groups of 10 headless mosquitoes were transferred to a 1.5-ml tube. Midguts from four groups of 10 mosquitoes, previously surface-sterilized, were dissected in a drop of sterile PBS and transferred to a 1.5-ml tube containing 50 µl of sterile PBS. All samples were flash frozen in an ethanol/dry-ice bath and stored at -80°C until RNA extraction.

Midguts and headless mosquitoes were homogenized in 1ml of Trizol with Lysing Matrix D Fast Prep tubes in the FastPrep 24 (MP Bio), followed by RNA extraction using the PureLink RNA Mini kit with on-column DNase treatment (ThermoFisher). RNA-Seq Libraries were prepared using the Universal Plus mRNA-Seq Library prep kit (Tecan Genomics) incorporating unique dual indexes. Libraries were assessed for quality by High Sensitivity D5000 ScreenTape on the 4200 TapeStation (Agilent Technologies). Quantification was performed with NuQuant reagent and by Qubit High Sensitivity DNA assay, on Qubit 4 and Qubit Flex Fluorometers (Tecan Genomics/ThermoFisher).

Libraries were diluted and an equimolar pool was prepared, according to the manufacturer’s protocol for appropriate sequencer. An Illumina iSeq Sequencer with iSeq100 i1 reagent V2 300 cycle kit was used for the final quality assessment of the library pool. For deep mRNA sequencing, a 300 cycle (2 x 150 bp) Illumina NextSeq 500 (High) run was performed. Gene coverage was about 40-50% with the average total reads at 24 million per sample. Three-to-four biological replicates were analyzed for each condition for a total of 15 sequencing libraries. RNA-seq data was analyzed with Partek Flow NGS Software as follows: pre-alignment QA/QC; alignment to *A. gambiae* Reference Index using STAR 2.7.8a; post-alignment QA/QC; quantification of gene counts to annotation model (Partek E/M); noise reduction filter by excluding features across samples where maximum ≤10; normalization of gene counts using Median Rario (DESeq2 only, Supplementary Data 2); and identification and comparison of differentially expressed genes with DESeq2. Reference Genome: NCBI: GCA_000005575.1_AgamP3. From the resulting gene lists, gene enrichment and pathway enrichment were performed within Partek Flow (Supplementary Fig. 1b). Genes of interest were validated by q-RTPCR. All sequence files and sample information have been deposited at NCBI Sequence Read Archive, NCBI BioProject: PRJNA1161683.

### A. gambiae infection with P. falciparum

Newly emerged adult female mosquitoes were allowed to feed on sugar solutions with up to 1 mM L-DOPA or dopamine for 4 d. On the fifth day, sugar meals were removed, and mosquitoes were starved for 5-6 h prior to offering a *Plasmodium falciparum* NF54-laced blood meal with gametocytemia 0.03%. Unfed mosquitoes were sorted, and blood-engorged mosquitoes were further incubated for 8 d at 27 °C for oocyst counting. Midguts from *Pf*-infected mosquitoes were dissected, stained with a 0.2% mercurochrome solution, and mounted on a slide with a coverslip gently pressed on top to facilitate oocyst visualization. Enumeration of oocysts was performed using a 20X objective under a Leica light microscope. For infections under aseptic conditions, the artificial glass feeder membrane-feeding process was conducted under conditions as sterile as possible to prevent antibiotic-treated mosquitoes from acquiring a bacterial infection ^98^.

### *In vitro* susceptibility of *P. falciparum* NF54 to L-DOPA

To determine *P. falciparum* toxicity to L-DOPA, solutions at concentration up to 2 mM were freshly prepared from a 5 mM L-DOPA stock. *P. falciparum* NF54 asexual and gametocyte stage cultures were dispensed into 96 well plates and triplicate wells were treated with vehicle or L-DOPA at concentrations of 0.25, 0.5, 1 and 2 mM for a total of 48 h. Blood smears were prepared after drug treatment and asexual parasitemia or gametocytemia was calculated microscopically from Giemsa-stained slides. The effect of L-DOPA exposure was determined by calculating % decrease in number of parasites as compared to vehicle control.

### Fungal challenge

*Cryptococcus neoformans* (strain H99-GFP) was cultured for 48 h in rich media at 30 °C with moderate shaking. Yeast cells were collected, washed twice with PBS, and adjusted to a final concentration of 1 x 10^8^ cells/ ml. In parallel, newly emerged adult female mosquitoes were allowed to feed with sugar solutions in the presence or absence of 1 mM L-DOPA for 4 d. Using a nanoinjector, adult mosquitoes were injected intrathoracically with 69 µl of the fungal suspension and then incubated at 19°C. Given that 1 mM L-DOPA triggers *C. neoformans* melanization enhancing fungal fitness against biotic and abiotic stressors ^119^, to assess antifungal response in *Anopheles* after infection, mosquitoes were surface sterilized 24 h post injection in 70% ethanol for 3 min and rinsed twice in PBS for 2 min each time. Individual mosquitoes were transferred to 1.5-ml tubes containing 100 µl PBS and homogenized for 20 s using a sterile pestle. Two serial dilutions of each sample were plated in Bird Seed agar (HiMedia Laboratories, PA), a selective media for isolation and differentiation of *Cryptococcus* spp, and incubated at 30°C for 7 d, after which the number of fungal colony-forming units (CFUs) were counted. Mosquito abdomens were also dissected to visualize the presence of live (GPF-positive) or melanized and dead (GFP- negative) *C. neoformans* yeasts, and images were captured using a Leica DM 2500 microscope.

### Cuticle pigmentation

Newly emerged adult female mosquitoes were allowed to feed with L-DOPA solutions for up to 14 d as described above. For photography, mosquitoes were cold anesthetized, placed ventral side up on a slide under a dissection microscope, and imaged with an Iphone X under constant exposure and lighting conditions. Image analysis and pigment measurement were done as described previously ^120^. Using the Fiji software ^121^, RGB color images were converted to 8-bit color, entire abdomens were selected using a freehand selection tool, and mean gray values were analyzed with the Measure tool. Pure black corresponds to 0 gray value whereas pure white equals 255.

### Transmission Electron Microscopy

Newly emerged adult mosquitoes were allowed to feed with sucrose solutions, with and without supplementation of 1 mM L-DOPA for 7 d. Groups of 10 midlegs from each condition were dissected on a glass petri dish. To reduce hydrophobicity, legs were rinsed once in 70% ethanol, twice in PBS for 1 min each step, and then incubated overnight at 4°C in fixing solution containing 2% glutaraldehyde, 0.1 M sodium cacodylate (pH 7.4) and 2% paraformaldehyde in distilled water. After buffer rinse, samples were postfixed in 1% osmium tetroxide, 0.8% potassium ferrocyanide in 0.1 M sodium cacodylate for at least one hour (no more than two) on ice in the dark. After osmium, samples were rinsed in 100 mM maleate buffer, followed by uranyl acetate (2%) in 100 mM maleate (0.22 µm filtered, 1 h dark), dehydrated in a graded series of ethanol and embedded in a SPURR’s/Eponate 12 (Ted Pella) resin. Samples were polymerized at 60C overnight. Thin sections, 60 to 90 nm, were cut with a diamond knife on the Reichert-Jung Ultracut E ultramicrotome and picked up with 2x1 mm formvar copper slot grids. To obtain consistent comparable segments across samples, femur cross-sections were obtained by conducting three sequential cuts at 200 nm depth into the femur from the direction of the tibia. Grids were stained with 2% uranyl acetate in 50% methanol followed by lead citrate and observed with a ThermoFisher Talos L120C at 120 kV. Images were captured with a ThermoFisher Ceta CCD (16 megapixel CMOS, 16-bit).

To account for differences in sequential cuts heterogeneity, at least three but no more than five representative images per cut were chosen per femur, each image captured ∼18 µm of total cuticle length. A minimum of 3 measurements of total cuticle and endocuticle per image were taken at the same point and exocuticle was calculated by subtracting these values. Areas of the cuticle containing structural modifications were excluded from measurements.

### Quantification of ^13^C L-DOPA Melanin by Solid-State NMR

Newly emerged adult mosquitoes were allowed to feed with sucrose solutions, with and without supplementation with 1 mM of stable-isotope enriched ^13^C L-DOPA (RING-^13^C6, 99%, CLM-1007- 0, Cambridge Isotope Laboratories, MA, USA) for 7 d. Next, mosquitoes were sorted by sex, collected in 15-ml Falcon tubes, and lyophilized for 3 d. Solid-state NMR measurements were conducted on 8-10 mg samples using a Varian (Agilent) DirectDrive2 spectrometer operating at a ^13^C frequency of 150 MHz and equipped with a 1.6-mm T3 HXY fast magic-angle spinning (fastMAS) probe spinning at 15.00 ± 0.02 kHz and a regulated temperature of 25 °C. ^13^C NMR spectra of the lyophilized mosquito samples were obtained with 90° pulse lengths of 1.2 and 1.4 μs for ^1^H and ^13^C, respectively; 104-kHz heteronuclear decoupling using the small phase incremental alternation pulse sequence (SPINAL^122^) was applied during signal acquisition. Whereas a 10% linearly ramped ^1^H amplitude, 1-ms time for ^1^H-to-^13^C polarization transfer, and 3-s delay between acquisitions were used to acquire CPMAS spectra that identified the carbon-containing functional groups, a longer 50-s delay was used for direct polarization (DPMAS) experiments that generated spectra with quantitatively reliable signal intensities. The CPMAS and DPMAS spectral runs required 4 and 56 hours, respectively.

### Chitin measurements

The chitin content of adult mosquitoes post L-DOPA feeding for 7 d was determined based on the method of Lehmann and White (1975), ^123^ adapted as described previously used to quantify the chitin content of the cryptococcal cell wall material ^124^. Using 15-ml tubes, 60 and 80 cold-anesthetized female and male mosquitoes, respectively, were freeze-dried, and lyophilized for 2- 3 d. Mosquitoes were transferred to 1.5-ml microcentrifuge tubes, weighted (typically 14 and 11 mg, respectively), and homogenized in 200 µl of distilled water using a battery-operated homogenizer (3 20-sec cycles) and a horned sonicator at half amplitude (2 30-sec cycles). To solubilize and alkali-extract the samples, homogenates were centrifuged at 21,000 *g* for 5 min at room temperature and the pellet of each sample was resuspended in an equal volume (200 µl) of 3% (w/v) of sodium dodecyl sulfate (SDS). The samples were then incubated at 100 °C for 15 min and centrifuged again for 5 min after cooling. Next, after each pellet was washed twice with 200 µl distilled water, it was resuspended in 150 µl 2.1 M KOH. Samples were incubated at 80 °C for 90 min and cooled down to room temperature. After 400 µl 75% ethanol was added to each sample, the sample was mixed and incubated on ice for 15 min. Sixty µl of a Celite 545 (Sigma-Aldrich) suspension (1g of Celite 545 in 12.5 ml 75% ethanol, rested for 2 min) was then added to each sample and centrifuged. The resultant cell matter was washed thoroughly once with 40% ethanol, twice with PBS (pH 7), twice with distilled water, and resuspended in distilled water to a final concentration of 10 mg ml ^-1^. To estimate the total amount of glucosamine stemming from chitin plus chitosan, one 100 µl aliquot (equivalent to 1 mg dry weight) was transferred into a 2 ml self-locking microcentrifuge tube and subjected to deacetylation with 100 µl 1 M HCl in a heating block at 110 °C for 2 h. As a blank, one 100 µl of milliQ water was used instead of the mosquito suspension. In a fume hood, sample deamination was achieved by adding 400 µl 2.5% NaNO2, then mixed and incubated for 15 min at room temperature to allow nitrogen oxides to dissipate. After that, 200 µl of 12.5% ammonium sulfate was slowly added to each sample, vortexed, and let stand at room temperature for 5 min. Measurement of hexosamines (amino sugars) from the deacetylated samples was done by treatment with 3-methyl-2-benzothiazolone hydrazine hydrochloride (MBTH, Sigma-Aldrich) under mildly acidic conditions. A 200 μl aliquot from each sample was transferred to a 96-well plate and the absorbance at 650nm was recorded. Concentrations were determined by comparison to a standard curve prepared from a 1mM GlcNAc stock (Sigma-Aldrich) and ranging from 0 to 100 nM.

### Tarsal contact bioassay

A susceptibility test against deltamethrin (0.05%) was done following the WHO protocol ^125^. The choice of deltamethrin, a pyrethroid insecticide, was justified by being one of the only two insecticide classes approved for use on insecticide-treated bed nets (ITNs). Using an aspirator 22-25 *A. gambiae* female mosquitoes aged 2-5 d and previously allowed to feed on sucrose with or without 1 mM L-DOPA supplementation for 4 d were introduced into two WHO holding tubes (one test and one control) that contained untreated papers. Mosquitoes were allowed to settle inside the tube for 15 min in an upright position, then with a quick and gentle tap mosquitoes were transferred into the exposure tubes containing the insecticide-impregnated papers. Following one-hour exposure at room temperature, mosquitoes were transferred back into holding tubes, provided with a cotton pad moistened with 10 % sugar, and incubated at 27 °C. The number of dead mosquitoes at 24 h was recorded. Control tubes contained oil-treated paper.

### Electron Paramagnetic Resonance (EPR) Spectroscopy

Acid-resistant material attributable to melanin was isolated from *A. gambiae* mosquitoes fed with and without 1 mM L-DOPA supplementation for 7 d following the protocol described previously ^126^. *Aedes aegypti* (mutant strain Yellow) and fungal eumelanin from *C. neoformans* were used as negative and positive controls, respectively. EPR measurements were conducted at the Caltech EPR Facility using a Bruker EMX X-band continuous-wave (CW) EPR spectrometer equipped with a Bruker SHQE resonator. Samples were measured at room temperature under non-saturating microwave power (4.4 µW) using a field modulation amplitude of 1 Gauss and conversion time of 5 ms.

### Infrared thermography for mosquito bodies

L-DOPA fed mosquitoes for 7 d were cold anesthetized, clustered into groups of ten on top of a cooling block set at 0°C and arranged from left to right for increasing L-DOPA concentrations for up to 1 mM. Basal mosquito temperatures were recorded in dark conditions using an infrared (IR) camera (FLIR Systems, Wilsonville, OR) mounted 40 cm above the cooling block setup on a benchtop in a temperature-controlled room (22 ± 5°C, 50% relative humidity). Irradiations were conducted for 4 min using a white LED lamp (250 W, 1850 Lumens, PAR38HO-E26-19W-4000K- 25°, Green Creative) mounted 40 cm above the cooling block and resulting in an average luminance of ∼415 LUX. The optimal LED bulb positioning for homogeneous light exposure was calculated based on the light intensity distribution using graph paper and a light meter (**Supplementary** Fig. 5).

### Fecundity, egg hatching rate and life span

For the fecundity assay, newly emerged adult female *A. gambiae* were allowed to feed on a sugar solution supplemented with 1mM L-DOPA for 7 d, whereas the controls were fed 1.5% sucrose without L-DOPA. Mosquitoes were allowed to blood-feed on mice for 20 min and then anesthetized on ice; all non-engorged mosquitoes were discarded. At 3 d post blood meal, individual mosquitoes were transferred to a 50-ml Falcon tube containing a paper cup in the bottom halfway submerged into clean distilled water for oviposition. The number of eggs laid by each mosquito was recorded for up to 4 d under a light microscope. Female mosquitoes that died before oviposition were not included in the study. After each count, adult mosquitoes were removed from the vials, eggs were completely submerged into larval rearing water, and vials were incubated for 3-4 d at 27°C and 30°C, making sure that the paper cups remained moistened. The first and second instar larvae were counted under a light microscope.

The life span of *A. gambiae* females during continued exposure to a sugar meal supplemented with 1 mM L-DOPA was measured by placing newly emerged mosquitoes into paper cups with a sugar solution that was freshly prepared every 2-3 d. Dead mosquitoes were recorded and removed daily until no live mosquitoes remained. For this experimental setting, control and 1mM L-DOPA cups were directly incubated inside the reach-in incubator. Survival analysis represents pooled data from three biological replicates of 35 mosquitoes each.

### Dopamine measurements

Dopamine quantification was conducted from whole female mosquitoes post L-DOPA feeding and from bacterial supernatants.

### Sample collection

For mosquito samples, newly emerged adult mosquitoes were allowed to feed with sugar solutions supplemented with L-DOPA at three different concentrations (0.2 mM, 0.6 mM, and 1 mM) for 4 d. At the assayed time, five whole female mosquitoes were collected in 1.5-ml brown tubes and flash-frozen in an ethanol-dry ice bath. Mosquito supernatants containing both hemolymph and cuticle dopamine were obtained by adding 200 µl ice-cold 0.2 M perchloric acid to each tube, homogenizing for 20 s on ice, and centrifuging at 20,000 g for 10 min at 4 °C^127^. Supernatants (∼100 µl) were transferred to a new brown 1.5-ml tube and kept at -80 °C until further processing was conducted. For bacterial samples, a single colony of each bacterial strain, previously shown to induce fungal melanization, was grown in LB broth at 30 °C for 96 h. Filter sterilized, cell-free culture supernatants were acidified to pH 4 with 1 M HCl, fractionated by FPLC using an AKTA Pure liquid chromatograph with a UV-900 monitor to measure absorbance at 214 nm (Amersham Biosciences) and a C18 reverse-phase column (Phenomenex Luna 5 µm, 100 Å, LC Column 150 x 4.6 mm) using an isocratic mobile phase of 25 mM potassium phosphate buffer (pH 3.1) at a flow rate of 1.0 ml/min ^53^. Fractions of 200 µl were collected into a new brown 1.5- ml tube and kept at -80 °C until further processing was conducted; a 50 mM dopamine (Sigma-Aldrich) solution was used as standard.

### Sample preparation and derivatization

Dopamine standards were prepared in 10 µg/mL sodium metabisulfite in water. Samples were derivatized as described previously, with the following adjustments: 20 µL of standards and samples were transferred to an extraction plate; 10 µL of 100 mM sodium tetraborate in water and 5 µL of a cold internal standard solution consisting of 500 ng/mL dopamine-d3 in acetonitrile (v/v) were added to each sample and mixed for 1 min on plate shaker at 1,600 rpm; 10 µL of 2% benzoyl chloride in acetonitrile (v/v) was then added to each sample and the plate was mixed for 2 min at 1,600 rpm. 10µL of 1% formic acid in water (v/v) was added to quench the reaction. 1µL of each sample was injected for analysis.

### Sample analysis

Samples were analyzed using an Agilent 6540 QTOF with a Jet Stream Electrospray Ionization Source (ESI) and an Agilent 1290 UHPLC (Agilent Technologies, Santa Clara, CA). Solvents were 10 mM ammonium formate in water (A) and acetonitrile (B). The samples were kept at 10°C during analysis. Chromatographic separation was achieved over 3.0 minutes using a Phenomenex Luna Omega 2.1 x 100mm, 1.6µm, C18 column at 40°C with a binary gradient starting with 10% B. The LC gradient was as follows: 0–1.2 min, 10% - 95% B; 1.2 - 2.0 min, 95% B; 2.01, 10% B. The flow rate was 400 µl min^−1^. Mass spectral acquisition was performed using full scan MS from *m/z* 100 to 1000 with the following source conditions: Drying and Sheath Gas temperatures of 350°C and 400°C, respectively; both gas flows at 12L/min; Nebulizer- 45psig, VCap, Nozzle and Fragmenter voltages at 3000 V, 600 V and 100 V, respectively. Proton adducts of the triple benzoylated dopamine and dopamine-d3 (*m/z* 466.1649 and 469.1837, respectively) were extracted within a +/- 20 ppm window. Analyte peak areas were determined from the extracted ion chromatograms and concentrations were calculated by linear regression analysis using Agilent Masshunter Quantitative Analysis Software (B.08.00). Limit of quantification was 1.0 ng/ml.

### Plate confrontation assays

As a proxy for the ability to synthesize catecholamines from bacteria isolated from the midgut of wild-caught *An. aranbiensis* and *Ae. aegypti* adult mosquitoes as well as an environmental source relevant to mosquito aquatic stages, we monitored Cryptococcal melanization as previously reported ^53^.

### Statistical Analysis

Data were graphed and analyzed for statistical significance using GraphPad Prism 10 (version 10.0.1) software (GraphPad Software, LLC). Particular tests used for experiments are indicated in the legend of each respective figure.

### Data availability

All data generated or analyzed during this study are included in this published article (and its Supplementary files). The datasets presented in this manuscript are publicly available Figshare data repository at 10.6084/m9.figshare.27122907 and 10.6084/m9.figshare.28063112. The RNAseq data was submitted to the Sequence Read Archive (SRA) database (PRJNA1161683).

## Acknowledgments

We would like to thank the Parasitology and Insectary Core Facilities at the Johns Hopkins Malaria Research Institute and Bloomberg Philanthropies for generous supplemental funding and core infrastructure that supported this research. We are grateful to the perseverance and loving memory of Ricardo Perez-Dulzaides for troubleshooting plate confrontation assays. Thank you to Diego Giraldo from Conor McMeniman lab for providing Aedes aegypti yellow mutant strain. RNA extraction, library preparation, iSeq run, qRT-PCR, NCBI SRA submission and analysis were performed by the JHSPH Genomic Analysis and Sequencing Core. Illumina NextSeq 500 run was performed at the Johns Hopkins Single Cell & Transcriptomics Core by Linda Orzolek and team. NMR measurements were conducted at the City College of New York Solid-state NMR Core Facility. EPR measurements were conducted by Paul H. Oyala at the Caltech EPR facility in the Division of Chemistry and Chemical Engineering (CCE).

## Author contributions

E.C.: conceptualization, methodology, validation, formal analysis, investigation, project administration, funding acquisition, writing—original draft, writing-review and editing, visualization. Y.D., C.C., R.J.B.C., R.G.S., Y.A-R., D.F.Q.S., E.J., I.H., J.A.P-M., M.DP., A.D., A.J., B.S., G.M., A.T.: methodology, validation, formal analysis, investigation, writing— writing-review and editing, visualization. N.A.B.: Resources, writing-review, and editing. R.E.S.: Resources, supervision, writing-review and editing, and funding acquisition. D.G.: Resources, supervision, validation, writing-review, and editing. A.C.: conceptualization, supervision, project administration, resources, supervision, writing-review and editing, and funding acquisition.

## Funding Acknowledgements

This work was supported in part by the National Institutes of Health grant number R01-AI171093 and JHMRI Pilot Grant US123 to A.C. E.C was also supported by the JHMRI Postdoctoral Fellowship, JHU COVID-bridge award 1316000001, and JHMRI Pilot Grant US139.

---

**Supplementary Data 1. Differentially Expressed Genes between 1mM L-DOPA-fed and sucrose-fed *A. gambiae* female mosquitoes.** 10.6084/m9.figshare.27122907

**Supplementary Data 2.** DESeq2 Median Ratio normalization for 902 differentially expressed genes in *A. gambiae* whole female mosquito post dietary L-DOPA. 10.6084/m9.figshare.28063112

**Supplementary Table 1.** Detection of melanin precursors released by diverse bacterial species.

**Supplementary** Fig 1**. Partek Flow analysis. A)** Principal component analysis (PCA) of 4 L-DOPA-fed mosquito groups and 3 sugar-fed controls. Blue spheres represent control replicates; red spheres represent L-DOPA-fed replicates. **B)** Top 20 KEGG Pathway enrichment analysis of transcriptional data.

**Supplementary** Fig 2. qRT-PCR analysis of mRNA abundance in whole mosquitoes from critical genes related to this study [phenoloxidase 5 (PPO5), 3,4-dihydroxyphenylacetaldehyde (DOPAL) synthase, thioester-containing protein 1 (TEP1), propionyl-CoA synthetase (ProCoA). Data represent the mean ± SE of three or more independent replicates, analyzed using unpaired by one-way ANOVA and multiple comparison test to determine gene expression relative to housekeeping S7 gene. NS, not significant

**Supplementary** Fig 3**. L-DOPA supplementation also enhances cuticular melanization in *Anopheles gambiae* male mosquitoes. A)** ^13^C DPMAS NMR spectra of ^13^C-L-DOPA fed mosquitoes display pigment peaks (green arrows) with greater relative intensities than their controls. Spectra were normalized to the tallest peak (lipid peak at ∼30 ppm). **B)** *Top panel,* TEM micrographs displaying distinct electron-dense bands within the endocuticle of L-DOPA-fed mosquitoes not seen in those of control. Scale bar, 500 nm. *Lower panel,* Measurements of cuticular thickness from legs of male mosquitoes fed with a 1 mM L-DOPA supplemented sugar meal show reduced exocuticle in comparison to controls (sugar-fed). NS, not significant; \*\*\*\**p* < 0.0001; Two-way analysis of variance (ANOVA) with Tukey’s multiple comparisons test.

**Supplementary** Fig 4**. Mapping white LED intensity distribution on the cooling block and mosquitoes’ temperature variance. A)** Apparent temperature of group of 20 mosquitoes fed with up to 1 mM L-DOPA demonstrated that 1 mM L-DOPA-fed mosquitoes exhibited a warmer temperature than sugar-fed controls (Trial 1). **B)** Apparent temperature of group of 20 mosquitoes fed with 1 mM L-DOPA consistently demonstrated that 1 mM L-DOPA-fed mosquitoes exhibited a warmer temperature that sugar-fed controls (Trial 2). **C)** Apparent temperature of individual mosquitoes (n = 6) fed with 1 mM L-DOPA consistently demonstrated that 1 mM L-DOPA-fed mosquitoes exhibited a warmer temperature that sugar-fed controls (Trial 3). **D)** Light intensity (LUX) distribution on sample platform showing that luminous flux decreases around the edge of the cooling block.

**Supplementary** Fig 5**. Gut microbiota can secrete dopamine.** Chromatographs of cell-free bacterial supernatants after growth for 24, 48, 72 and 96 h Luria Bertani media at 30°C. Overlapped dotted chromatograph corresponds to 1 mM spike dopamine. Red arrows show points at which samples were collected. The elution time was measured in minutes and the absorbance is measured at 214 nm in arbitrary units. For Fast Protein Liquid Chromatography (FPLC), we used a C18 Reverse-Phase column to observe the dopamine peaks. 25 mM Potassium Phosphate buffer (pH 3.1) was used as an isocratic mobile phase at a flow rate of 1 ml/min.

**Supplementary** Fig 6**. L-DOPA-sucrose solutions pH monitored over time.** Measurement of pH from sucrose solution while incubated at 27°C inside the mosquito chamber displays progressive acidification. Supplementation with L-DOPA accentuates this trend. Data corresponding to three independent measurements for all conditions. Bars correspond to mean with 95% confidence interval. Statistical analyses were performed using a Two-way analysis of variance (ANOVA) with Sidak’s multiple comparisons test.

**Table 1.**
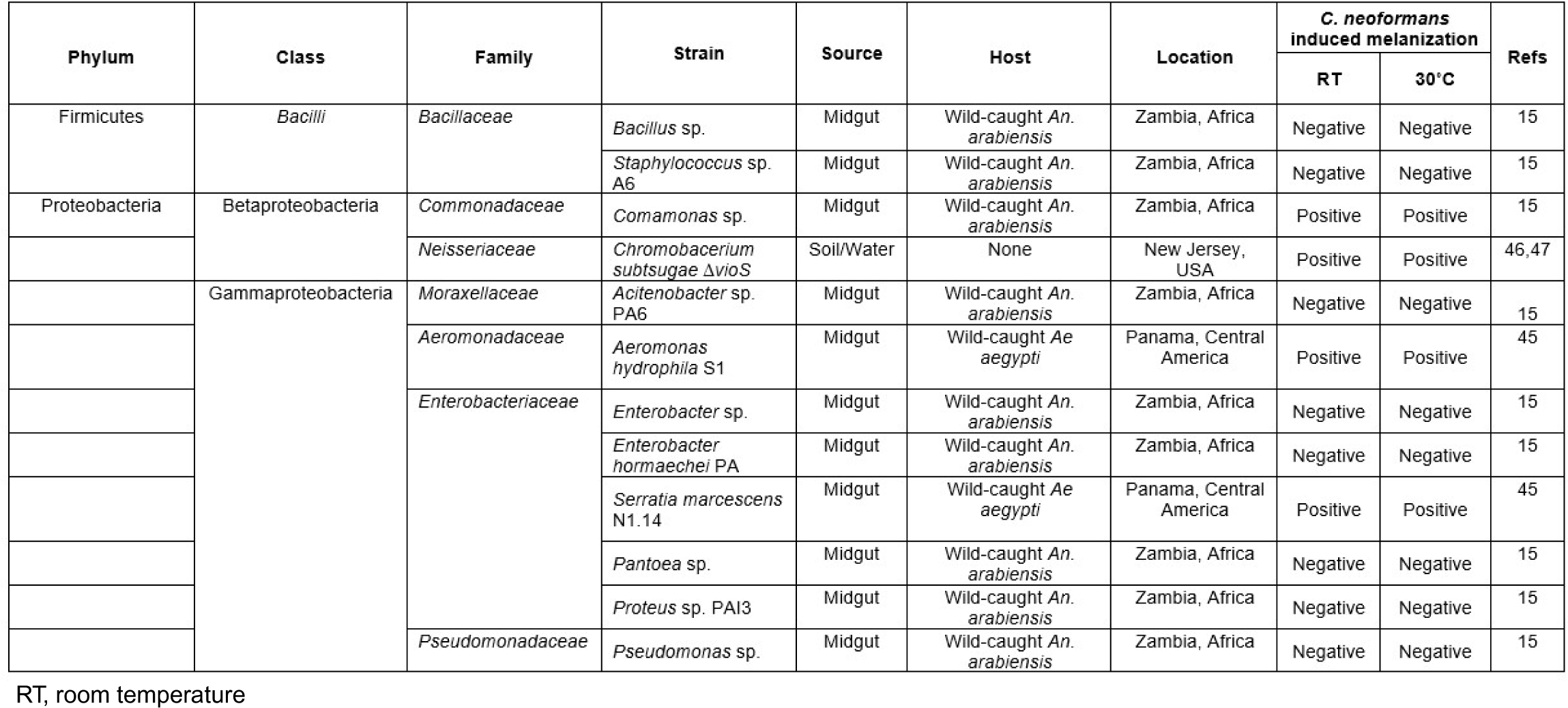
Detection of melanin precursors released by diverse bacterial species.

