## Supplemental Material for "Dietary L-3,4-dihydroxyphenylalanine (L-DOPA) augments cuticular melanization in *Anopheles* mosquitos while reducing their lifespan and malaria parasite burden"

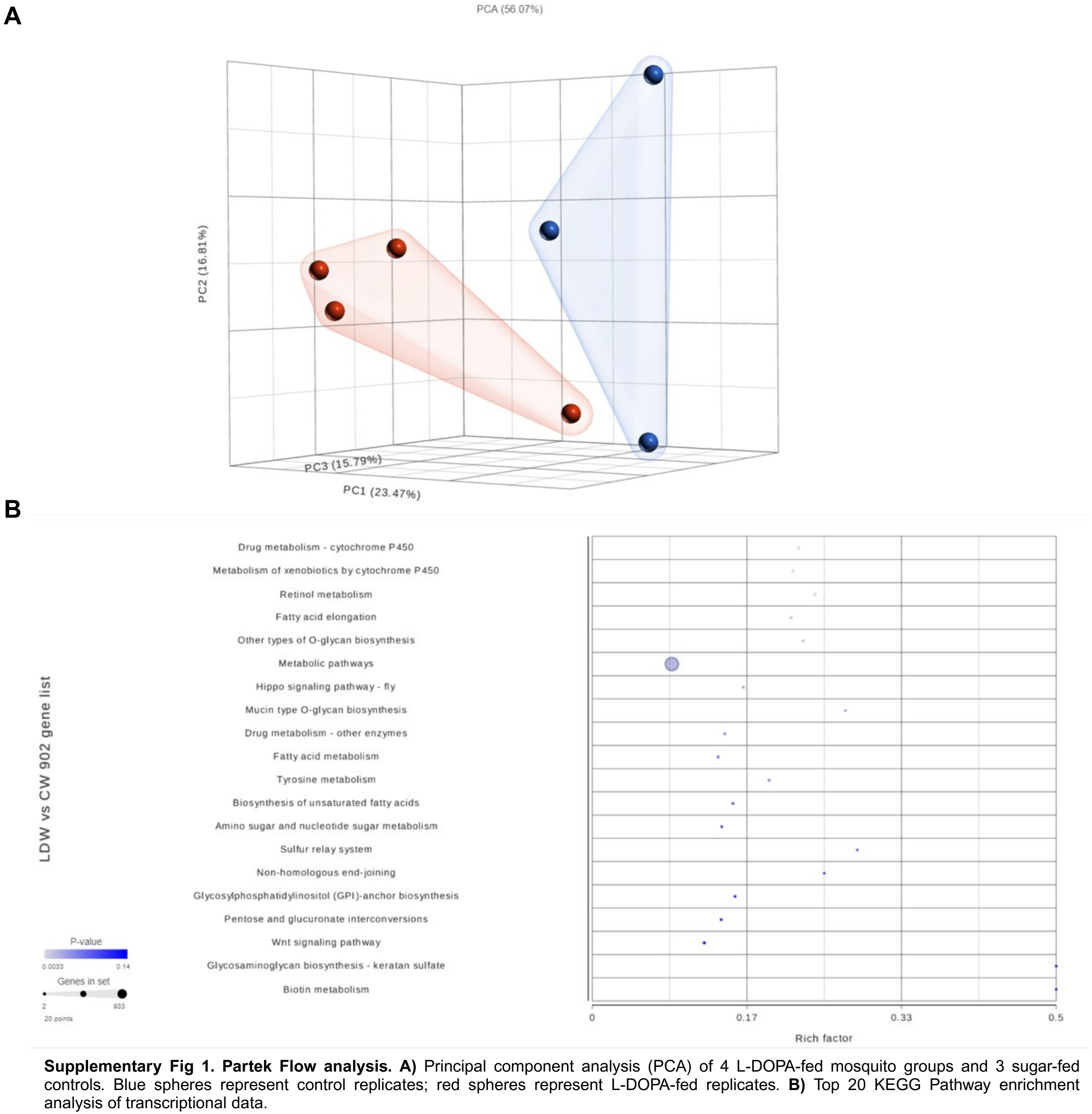


**Supplementary Fig 1. Partek Flow analysis. A)** Principal component analysis (PCA) of 4 L-DOPA-fed mosquito groups and 3 sugar-fed controls. Blue spheres represent control replicates; red spheres represent L-DOPA-fed replicates. **B)** Top 20 KEGG Pathway enrichment analysis of transcriptional data.

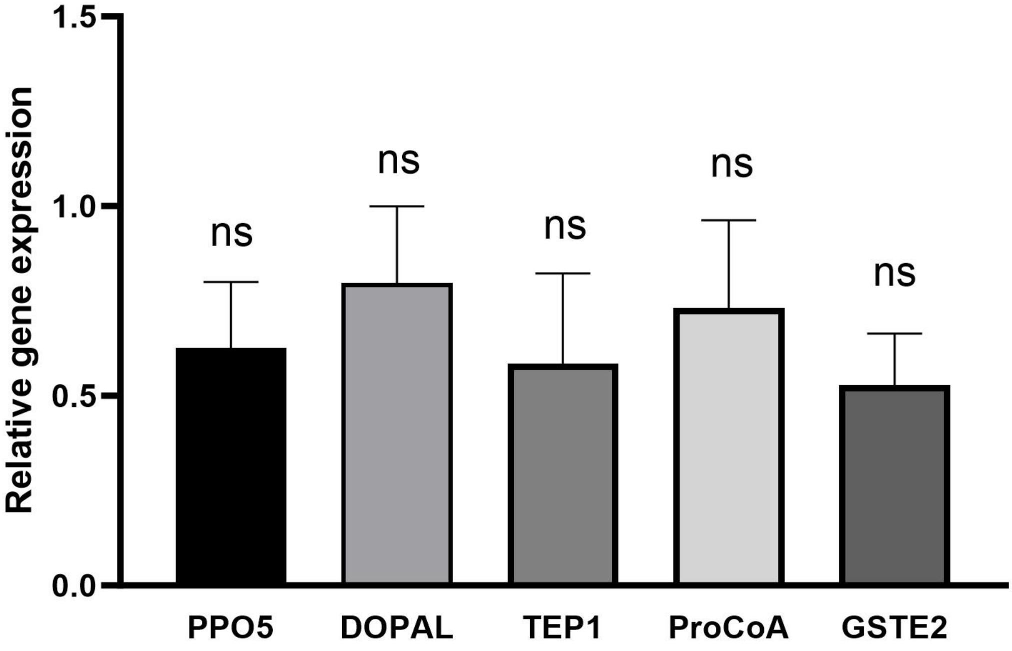


**Supplementary Fig 2.** qRT-PCR analysis of mRNA abundance in whole mosquitoes from critical genes related to this study [phenoloxidase 5 ( PPO5), 3,4-dihydroxyphenylacetaldehyde (DOPAL) synthase, thioester-containing protein 1 (TEP1), propionyl-CoA synthetase (ProCoA). Data represent the mean ± SE of three or more independent replicates, analyzed using unpaired by one-way ANOVA and multiple comparison test to determine gene expression relative to housekeeping S7 gene. NS, not significant

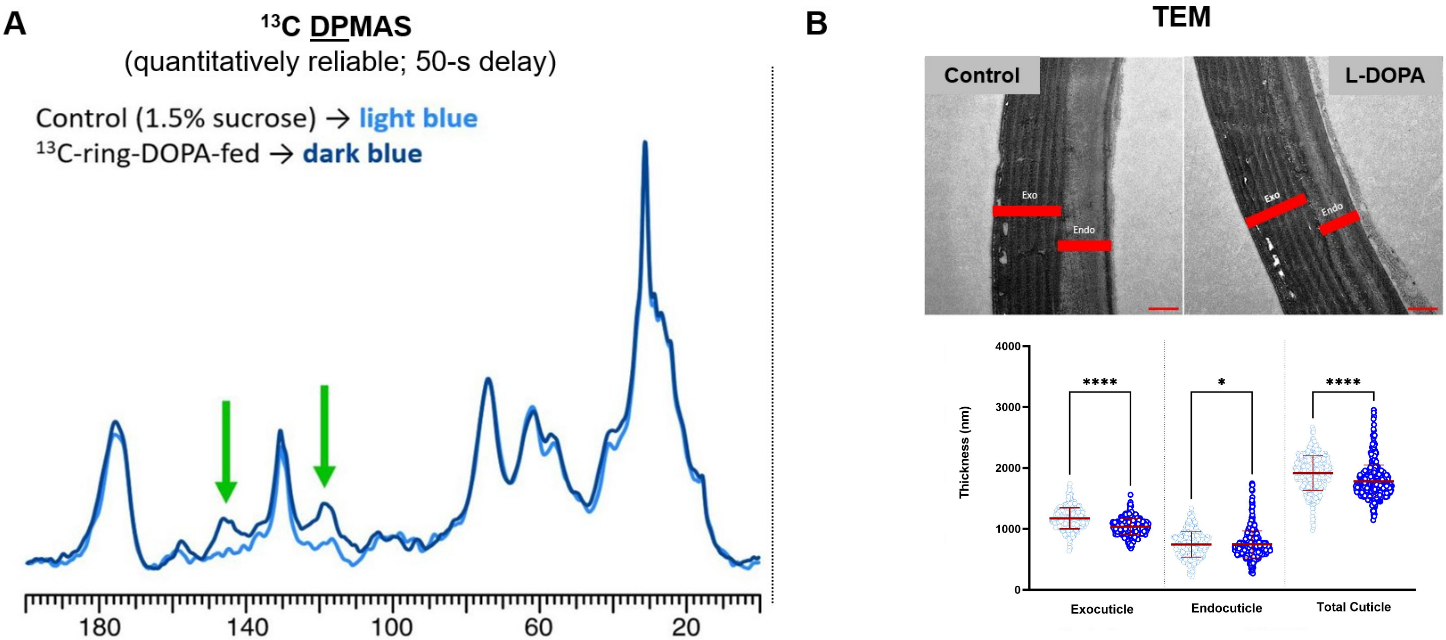


**Supplementary Fig 3. L-DOPA supplementation also enhances cuticular melanization in *Anopheles gambiae* male mosquitoes. A)** $^{13}$C DPMAS NMR spectra of $^{13}$C-L-DOPA fed mosquitoes display pigment peaks (green arrows) with greater relative intensities in comparison to their controls. Spectra were normalized to tallest peak (lipid peak at ~30 ppm). **B)** *Top panel,* TEM micrographs displaying distinct electron-dense bands within the endocuticle of L-DOPA-fed mosquitoes not seen in those of control. Scale bar, 500 nm. *Lower panel,* Measurements of cuticular thickness from legs of male mosquitoes fed with a 1 mM L-DOPA supplemented sugar meal show reduced exocuticle in comparison to controls (sugar-fed). Bar corresponds to mean with SD. Statistical analyses were performed using a Two-way analysis of variance (ANOVA) with Sidak's multiple comparisons test. **p*=0.0462, *****p*<0.0001; Two-way analysis of variance (ANOVA) with <0.0001.

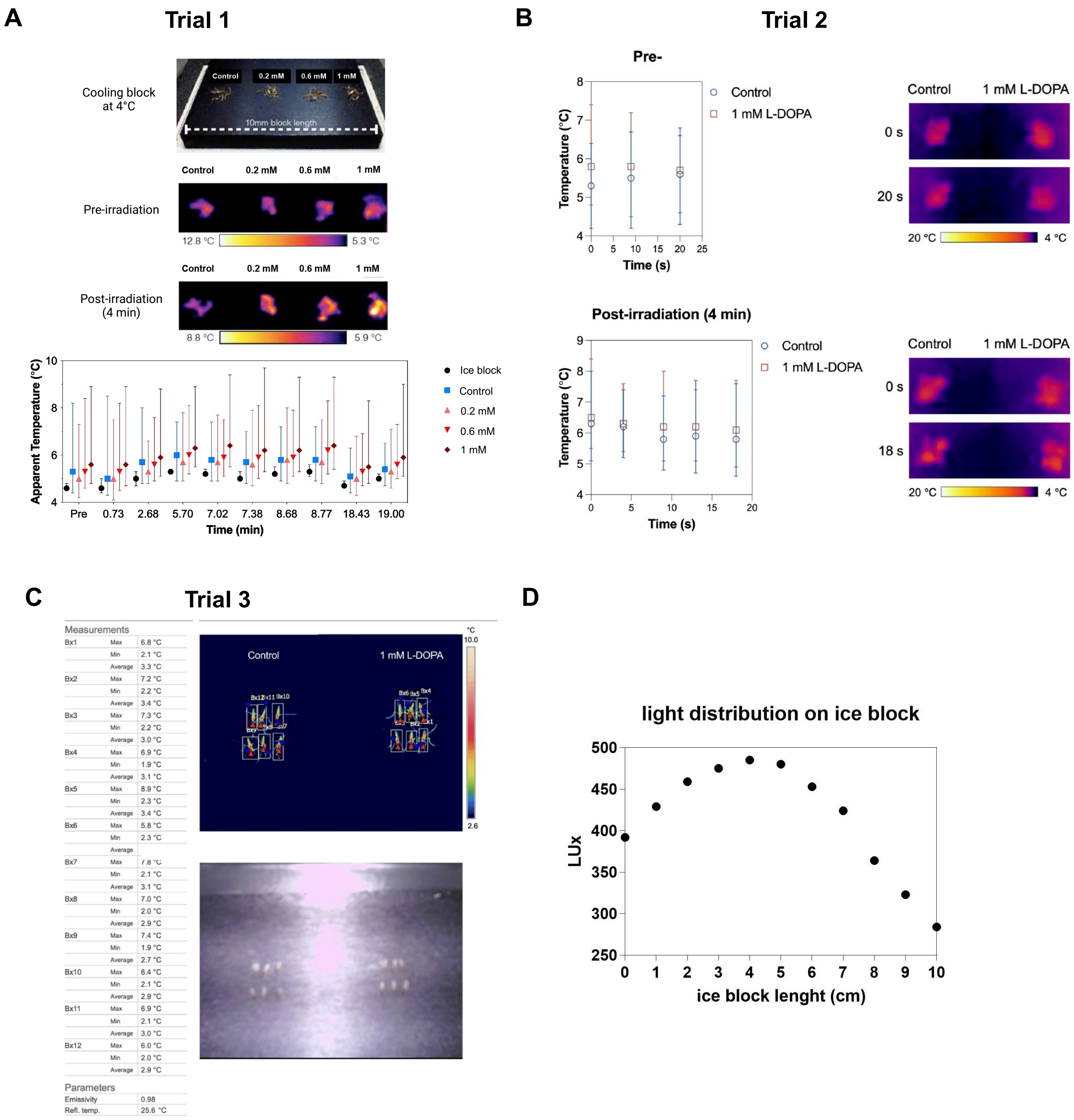


**Supplementary Fig 4. Mapping white LED intensity distribution on the cooling block and mosquitoes' temperature variance. A)** Apparent temperature of group of 20 mosquitoes fed with up to 1 mM L-DOPA demonstrated that 1 mM L-DOPA-fed mosquitoes exhibited a warmer temperature than sugar-fed controls (Trial 1). **B)** Apparent temperature of group of 20 mosquitoes fed with 1 mM L-DOPA consistently demonstrated that 1 mM L-DOPA-fed mosquitoes exhibited a warmer temperature that sugar-fed controls (Trial 2). **C)** Apparent temperature of individual mosquitoes (n = 6) fed with 1 mM L-DOPA consistently demonstrated that 1 mM L-DOPA-fed mosquitoes exhibited a warmer temperature that sugar-fed controls (Trial 3). **D)** Light intensity (LUX) distribution on sample platform showing that luminous flux decreases around the edge of the cooling block.

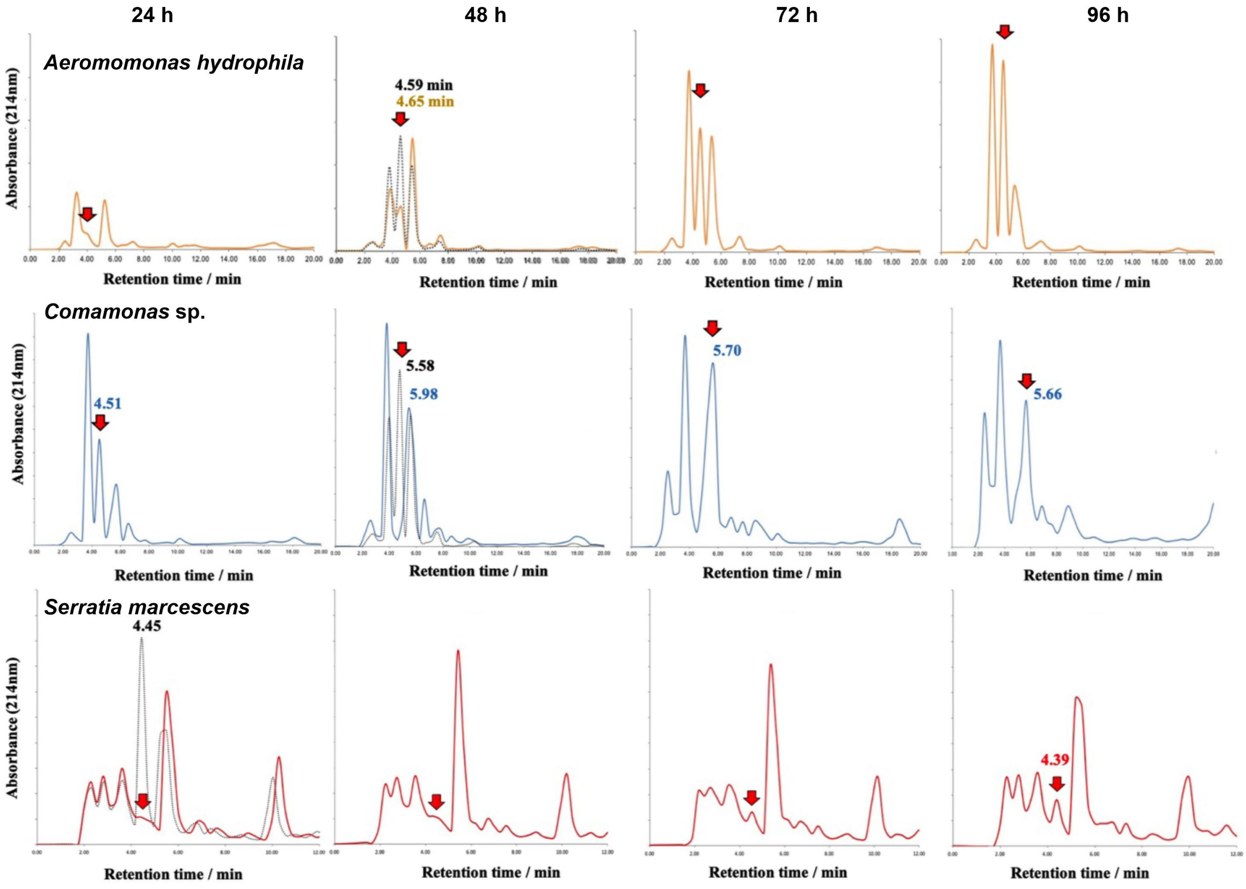


**Supplementary Fig 5. Gut microbiota can secrete dopamine.** Chromotographs of cell-free bacterial supernatants after growth for 24, 48, 72 and 96 h Luria Bertani media at 30$^{o}$C. Overlapped dotted chromatograph corresponds to 1 mM spike dopamine. Red arrows, points were samples were collected. The elution time was measured in minutes and the absorbance is measured at 214 nm in arbitrary unit. For Fast Protein Liquid Chromotography (FPLC), we used a C18 Reverse-Phase column to recovered dopamine peaks. 25 mM Potassium Phosphate buffer (pH 3.1) was used as isocratic mobile phase at a flow rate of 1 ml/ml.

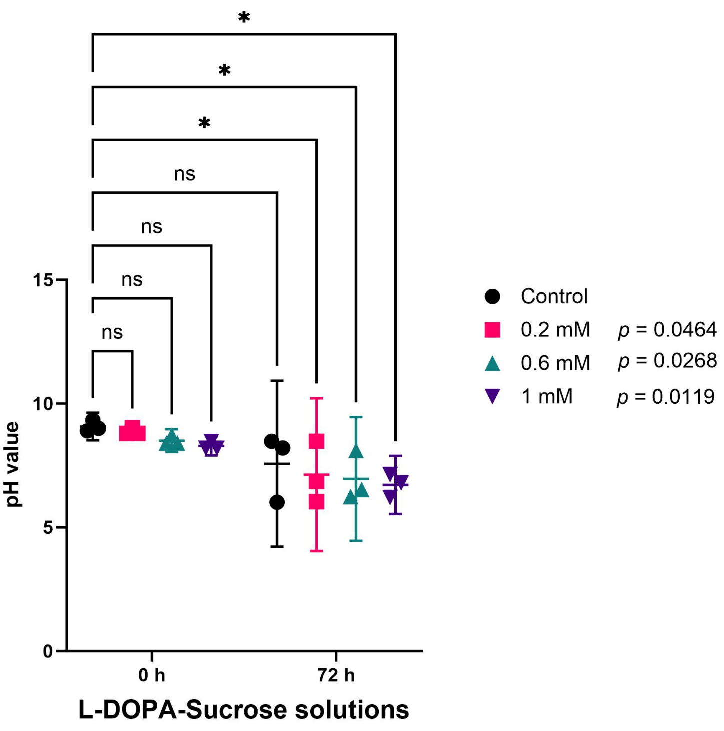


**Supplementary Fig 6. L-DOPA-sucrose solutions pH over time.** Measurement of pH from sucrose-solution while incubated at 27°C inside the mosquito chamber displays constant acidification. Supplementation with L-DOPA accentuates this reaction. Data corresponding to three independent measurements for all conditions. Bars correspond to mean with 95% CI. Statistical analyses were performed using a Two-way analysis of variance (ANOVA) with Sidak's multiple comparisons test.
